# Uncertainty-aware quantitative analysis of the structure and dynamics of T cell receptor repertoires

**DOI:** 10.64898/2026.06.28.735097

**Authors:** Simo Kitanovski, Kai Wollek, Daniel Hoffmann

## Abstract

Diversity and dynamics of immune cell receptor repertoires (IRRs) are two factors at the functional heart of adaptive immunity that together make IRRs difficult to grasp. Moreover, measurements are compounded by various sources of experimental noise. Here we propose a computational framework (*ClustIRR*) for uncertainty-aware quantitative analysis of IRR structure and dynamics. *ClustIRR* maps multiple IRRs across replicates, time points, or conditions onto a joint graph induced by immune receptor sequence similarity. It then detects *communities on the joint graph* (CJs). Based on CJs as reference structures across IRRs, *ClustIRR* then performs quantitative Bayesian analyses of differential CJ occupancy. Additionally, *ClustIRR* integrates single-cell gene expression data to link community expansion with transcriptional activation signatures. We demonstrate the capabilities of *ClustIRR* with the joint analysis of multiple T cell receptor repertoires in several example applications: (1) quantitative changes due to antigen challenge, (2) longitudinal dynamics during cancer immunotherapy, (3) V(D)J recombination biases in human vs murine repertoires that pre-adapt IRRs for pathogen responses. *ClustIRR* is freely available as open source software from the bioconductor repository.

## Introduction

Adaptive immunity relies on highly diverse immune receptor repertoires (IRRs) to recognize pathogens and malignant cells. Here we focus on T cell receptor (TCR) repertoires. TCR diversity arises through V(D)J recombination, generating an estimated 10^14^ possible receptor variants (***Murugan et al., 2012***). Within individuals, TCR repertoires are organized into clonotypes–subpopulations of cells sharing identical TCR sequences generated through antigen-driven clonal expansion. While clonally expanded T cells recognize identical antigens, accumulating evidence indicates that TCRs with non-identical but *similar* CDR3 sequences often recognize the same epitopes (***Glanville et al., 2017; Dash et al., 2017***). Such cross-reactivity has a number of consequences: for instance, even if clonotypes are not shared between individuals or samples, the respective repertoires may still recognize the same antigens. Conversely, the same TCR may recognize multiple antigens. In both directions cross-reactivity makes data interpretation more difficult.

High-throughput sequencing (HTS) of TCR repertoires (***Woodsworth et al., 2013; Heather et al., 2018; Pai and Satpathy, 2021***), and more recently single-cell TCR sequencing (***Han et al., 2014; Redmond et al., 2016***), enables precise clonotype enumeration at cellular resolution. To address sequence degeneracy, similarity-based clustering approaches group clonotypes into *communities* with putatively shared antigen specificity (***Glanville et al., 2017; Dash et al., 2017; Mayer-Blackwell et al., 2021; Zhang et al., 2020; Valkiers et al., 2021; Pogorelyy et al., 2019***). The architecture of TCR repertoires—defined by community composition and abundance—encodes critical functional information. For instance, successful cancer immunotherapy is associated with the expansion of tumor-antigen-specific communities, while viral exposure elicits cognate TCR communities. Consequently, the functional interpretation of TCR repertoires hinges on the ability to quantitatively track *community occupancy dynamics* across repertoires, rather than relying on static community identification within isolated samples.

However, achieving robust quantitative tracking requires to overcome significant hurdles rooted in the biological and technical nature of TCR repertoire data. First, quantifying changes at the clono-type level is confounded by vast diversity and limited sampling depth. Stochastic V(D)J recombination and experimental noise mean that identical clonotypes are rarely shared across repertoires, preventing direct detection of the same clone across conditions. Second, while coarse-graining by partitioning into communities mitigates clonotype sparsity, it introduces a structural challenge: communities must be defined *consistently* across samples to be comparable. Existing clustering tools typically operate on single repertoires in isolation, producing incompatible community labels that preclude longitudinal or cross-condition quantitative comparison.

Third, IRR sequencing data are inherently count-compositional, sparse, and noisy. These properties extend to community occupancy estimates. Standard differential abundance tests for TCR repertoires often focus on clonotypes and rely on qualitative comparisons based on point estimates (***Rudqvist et al., 2018***) or simple univariate approaches, such as beta-binomial counts regression (***Rytlewski et al., 2019; Mayer-Blackwell et al., 2021***) or Fisher’s exact test (***Balachandran et al., 2017***). Such approaches can mis-characterize uncertainty by overlooking key variance components, failing to distinguish biological expansion from stochastic fluctuation.

Bayesian hierarchical models offer a rigorous alternative, but these models have been rarely developed for differential abundance analysis in TCR repertoires. Existing Bayesian methods primarily focus on inferring latent clonotype frequencies from bulk sequencing read counts (***Puelma Touzel et al., 2020; Koraichi et al., 2022***). While these methods represent important advancements, they typically require biological replicates at each time point to quantify natural variability—a condition often unmet in clinical or longitudinal studies. Tools like *scCODA* (***Buettner et al., 2021***) or *tascCODA* (***Ostner et al., 2021***) address compositional bias in single-cell data but are designed for annotated cell types rather than sequence-defined TCR communities. They are also ideally suited for moderate dimensionality (typically < 100 categories), whereas TCR communities can number in the thousands. Consequently, there is currently no unified method that combines joint community definition across samples with probabilistic quantification for high-dimensional IRR data.

Here we introduce *ClustIRR*, a computational framework that addresses these interconnected challenges through model-based quantification of community occupancy in IRRs. While we focus on TCR repertoires, *ClustIRR* is equally applicable to B cell receptor (BCR) repertoires. Recognizing that robust quantification is contingent upon consistent community definitions, *ClustIRR* first constructs a *joint graph* integrating multiple repertoires across replicates, time points, or biological conditions. In this graph, nodes correspond to clonotypes, and edges reflect CDR3 sequence similarity based on pairwise alignment scores. *ClustIRR* employs community detection algorithms (e.g., Leiden (***Traag et al., 2019***), Louvain (***Blondel et al., 2008***)) to identify densely interconnected TCR groups suggesting shared antigen recognition. This process defines *Communities on the Joint graph* (CJs) that persist across repertoires.

To quantify dynamics, *ClustIRR* models cell counts per CJ across repertoires using a hierarchical Bayesian Dirichlet-Multinomial framework. By explicitly accounting for the count-compositional nature of the data, this approach quantifies both relative CJ occupancy and differential occupancy between repertoires, generating posterior distributions with 95% credible intervals (CIs). This enables researchers to distinguish true biological expansion from stochastic fluctuations. Additionally, *ClustIRR* integrates single-cell RNA sequencing (scRNA-seq) data, linking TCR CJ expansion with transcriptional states such as activation, exhaustion, and cell cycle progression. Finally, *ClustIRR* annotates CJs with antigen specificity through integration with public databases (VDJdb, TCR3d, McPAS) (***Shugay et al., 2018; Gowthaman and Pierce, 2019; Tickotsky et al., 2017***).

We demonstrate the capabilities of *ClustIRR* with three applications: (i) cross-condition repertoire comparison identifying antigen-specific CJs; (ii) longitudinal therapy monitoring revealing predictive biomarkers in cancer immunotherapy; and (iii) multi-replicate cohort analysis uncovering intrinsic V(D)J recombination biases between humans and mice. We further showcase scRNA-seq integration linking expansion of CJs with T cell activation signatures. Together, these applications demonstrate how *ClustIRR* enables quantitative, biologically meaningful assessment of TCR repertoire dynamics with rigorous uncertainty quantification. *ClustIRR* is available as an open-source R package from the Bioconductor repository.

## Results

### Detection of EBV- and MART1-antigen reactive T-cell communities from single cell datasets

To validate *ClustIRR*, we applied it to five single-cell TCR repertoires (Dataset 1, Methods), including three from healthy donors (C1, C2, C3) and two antigen-stimulated repertoires: Epstein-Barr virus (EBV; repertoire E) and MART1 (repertoire M). *ClustIRR* should show expansion of EBV and MART1 specific CJs, respectively, in the stimulated repertoires compared to repertoires from healthy donors.

*ClustIRR* constructed a joint graph comprising 89,710 nodes (T-cell clonotypes) connected by approximately 13 million edges, where edge weights reflected the average CDR3*αβ* sequence similarity scores between clonotype pairs (Methods). Application of the Leiden algorithm (Potts model, resolution parameter *r* = 1) identified 10,301 CJs. Among these, 3,038 were singletons, while the largest CJ contained 1,339 clonotypes representing 29,491 cells, 99.5% of which originated from the EBV-stimulated repertoire (E). Among non-singleton CJs, the maximum unweighted distance between any two nodes (i.e., the CJ diameter) was 2 (Supplementary Figure S1B). This indicates that the CJs are compact and well-connected, reflecting conserved CDR3 sequences. Importantly, connectivity within most CJs was driven by high similarity in either CDR3*α* or CDR3*β* sequences, but rarely both simultaneously (Supplementary Figure S1C). This pattern suggests that antigen specificity is primarily encoded by a single chain, permitting flexible pairing with diverse partner chains. Furthermore, this observation aligns with the principle that TCR chain pairing is nearly independent, which makes the convergent generation of identical *αβ* pairs highly improbable (***Dupic et al., 2019***).

To quantify repertoire overlap, *ClustIRR* computed pairwise cosine similarities (CS) using CJ occupancy vectors (cell counts per CJ) for each repertoire pair (Fig. 1A). Unstimulated controls had elevated inter-repertoire similarity from CS ≈ 0.5 between C3 and either of C1 and C2 to CS ≈ 0.9 between C1 and C2. In contrast, antigen-stimulated repertoires overlapped much less with controls (CS: 0.03–0.09 for E; 0.18–0.3 for M) and with each other (E vs. M: CS = 0.01).

**Figure 1.**
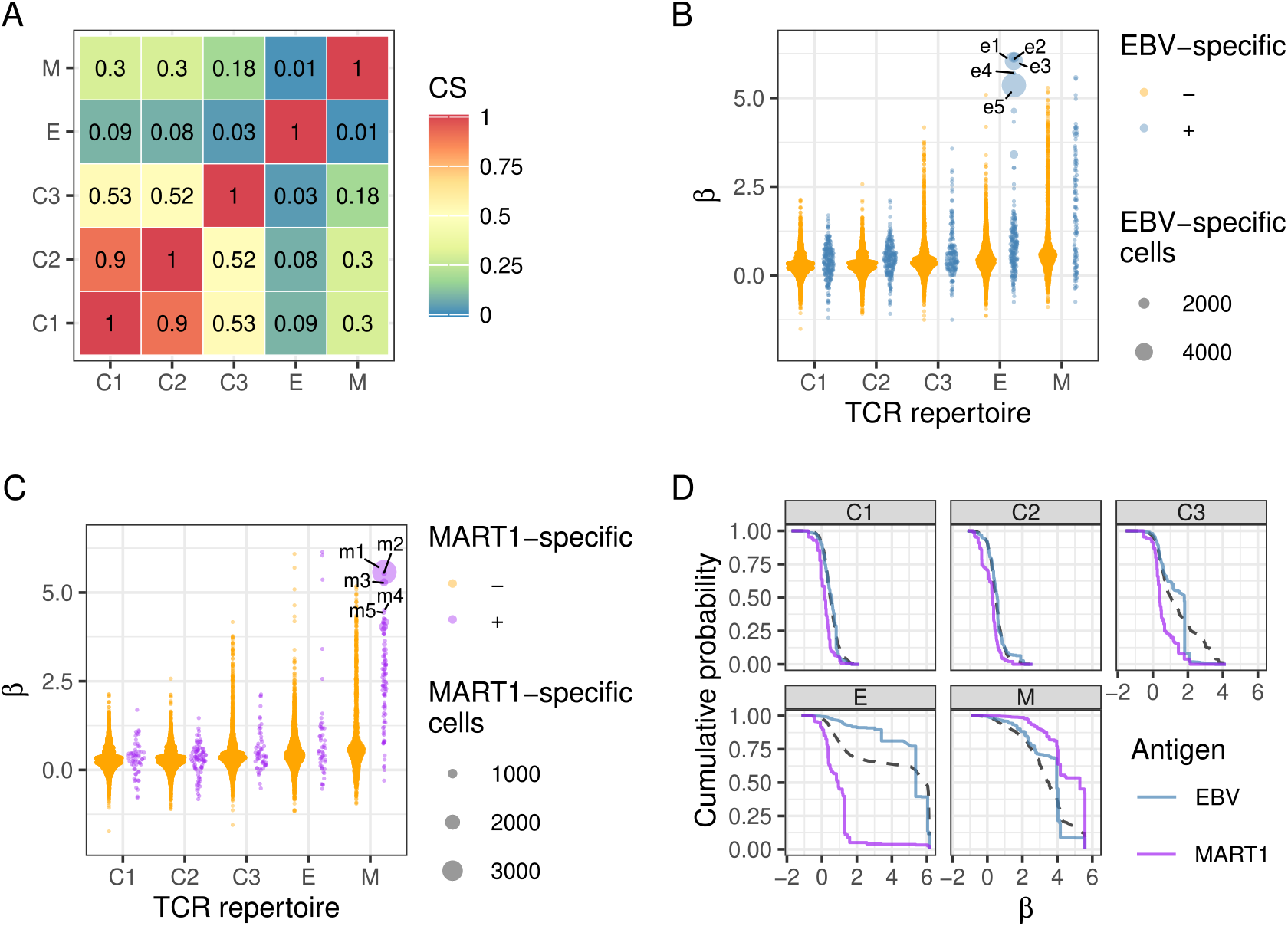
*ClustIRR* analysis of five TCR repertoires from Dataset 1. (A) Pairwise cosine similarity (CS) matrix between TCR repertoire’s CJ occupancy vectors (cell counts per CJ). Tile color intensity and labels indicate similarity values. (B) Mean *β* coefficients (y-axis) for 7,505 CJs (dots) across repertoires (x-axis). CJs categorized as EBV specific (+, blue) or non-EBV specific (–, orange). (C) Mean *β* coefficients (y-axis) for 7,505 CJs (dots) across repertoires (x-axis). CJs categorized as MART1 specific (+, purple) or non-MART1 specific (–, orange). Dot sizes indicate numbers of antigen specific T cells. Horizontal jitter width scales with point density. (D) Cumulative probability (y-axis) for all T cells (black) and T cells specific for EBV (blue) or MART1 (purple) across CJs ranked by *β* coefficients (x-axis) in each TCR repertoire (panels).

We quantified repertoire-specific CJ expansion using *ClustIRR*’s Bayesian model, which expresses the effect of repertoire *x* on CJ *i* occupancy by the coefficient *β*_*ix*_. The sign and magnitude of *β*_*ix*_ values indicate expansion (*β*_*ix*_> 0) or contraction (*β*_*ix*_< 0) in repertoire *x* (Methods). Given that adaptive immunity responds to antigenic challenge through clonal expansion, we hypothesized that expanded CJs in repertoires E and M would be enriched for EBV- and MART1-specific clonotypes, respectively. Consistent with this, *β*_*ix*_ distributions for E and M had heavy tails with large positive values beyond 5, in contrast to healthy controls (Fig. 1B-C) with *β* coefficients below 5.

Leveraging known CDR3-antigen associations from VDJdb, we identified 218 and 86 CJs in E and M contained at least one EBV- or MART1-specific T cell clonotype, respectively. Notably, several such clonotypes had expanded dramatically to thousands of T cells (large blue (EBV) and purple (MART1) dots in Fig. 1B-C). CJs in repertoires E and M with large *β* coefficients were enriched in T cells with known specificity for EBV or MART1 antigens compared to random expectation based on CJ sizes (Fig. 1D). In contrast, control repertoires were depleted of T cells specific for these antigens. The prevalence of T cells specific for unrelated antigens (e.g., human cytomegalovirus, CMV) matched random expectations (Supplementary Figure S2).

We inspected the five CJs (e1 to e5; blue dots with labels in Fig. 1B) with the highest *β* coefficients in repertoire E. These CJs contained between 4,266 and 29,491 T cells distributed across 156 to 1,339 clonotypes (Supplementary Figure S4B). The majority of T cells (>99%) and clonotypes (>69%) originated from repertoire E. Four CJs (e1, e2, e3, and e5) contained many T cell clonotypes with CDR3 sequences specific for EBV antigens according to VDJdb, with the most expanded clonotypes recognizing the epitope GLCTLVAML (EBV protein BMLF1_280-288_) restricted by HLA-A*02 (Supplementary Figure S4A). This epitope matched the antigen used to stimulate repertoire E from the donor carrying HLA-A*02. These clonotypes also matched VDJdb-annotated variable and joining gene segments. CJ e4 was expanded in EBV and contained conserved CDR3*β* sequences; however, their specificity was not determined, which does not preclude EBV-antigen specificity. Notably, these CJs (e1–e5) were among the most compact CJs in our repertoires (Supplementary Figure S1B).

Similarly, we inspected five CJs (m1 to m5; purple dots with labels in Fig. 1C) with the highest *β* coefficients in repertoire M and had at least one MART1-specific clonotype. These contained between 110 and 4,332 T cells distributed across 10 to 523 clonotypes. The majority of T cells (>95%) and clonotypes (>70%) originated from repertoire M (Supplementary Figure S4B). The most expanded clonotypes recognized the HLA-A*02-restricted epitope ELAGIGILTV (MART1_26-35_; Supplementary Figure S4A), matching the antigen used to stimulate repertoire M. These clonotypes also matched VDJdb-annotated variable and joining gene segments. Additional CJs in M with large *β* coefficients lacked VDJdb annotated MART1 specificity (orange dots in Fig. 1C), likely due to database incompleteness. These CJs are prime candidates for *de novo* identification of MART1 specific TCRs. As with the EBV-specific CJs (e1–e5), these CJs were among the most compact in our data (Supplementary Figure S1B), perhaps with the exception of m2 and m5. The reduced compactness of the latter two reflected heterogeneous CDR3 sequences containing known MART1 recognition motifs with variable core residues (m2 and m5 sequence logos in Supplementary Figure S4A), which demonstrates *ClustIRR*’s ability to detect biologically meaningful TCR groups with variable sequence heterogeneity.

### Integration with scRNA-seq data reveals signatures of T-cell activation signatures in expanded CJs

The combination of single-cell gene expression (GEX) profiles with the TCR data allows for the identification of correlations between GEX and TCR space. Here, we compared the GEX profiles of CJs with their expansion scores (*β*) across repertoires C1, C2, C3, E, and M.

Clustering of GEX profiles with *scBubbletree* (***Kitanovski et al., 2024***) identified four robust, transcriptionally distinct cell clusters, visualized as *bubbles* at the tips of a dendrogram (Fig. 2A). Over 99% of cells from unstimulated samples (C1, C2, C3) were assigned to bubble 0 (Fig. 2B), which represented the largest cluster in the dataset (76,000 cells; 36.3% of total cells). This homogeneity underscores the transcriptional similarity of unstimulated cells. Notably, C1, C2, and C3 collectively contributed 99% of cells in bubble 0 (each sample accounting for 33% of the cluster), while samples E and M contributed only 0.2% and 0.1%, respectively (Fig. 2C). In contrast, stimulated samples E and M populated different clusters: 66% of cells from sample E and 88% from sample M went to bubbles 1 and 2, respectively, and the remaining cells from E and M co-clustered in bubble 3, with sample E contributing >85% of cells in this cluster (Fig. 2C). Hence, bubble 3 provides a unique opportunity to investigate cells with converging transcriptional profiles in response to stimulation by different antigens. The relative occupancy of expanded CJs e1–e5 and m1–m5 across bubbles mirrored overall E/M distributions, indicating no bubble specific enrichment.

**Figure 2.**
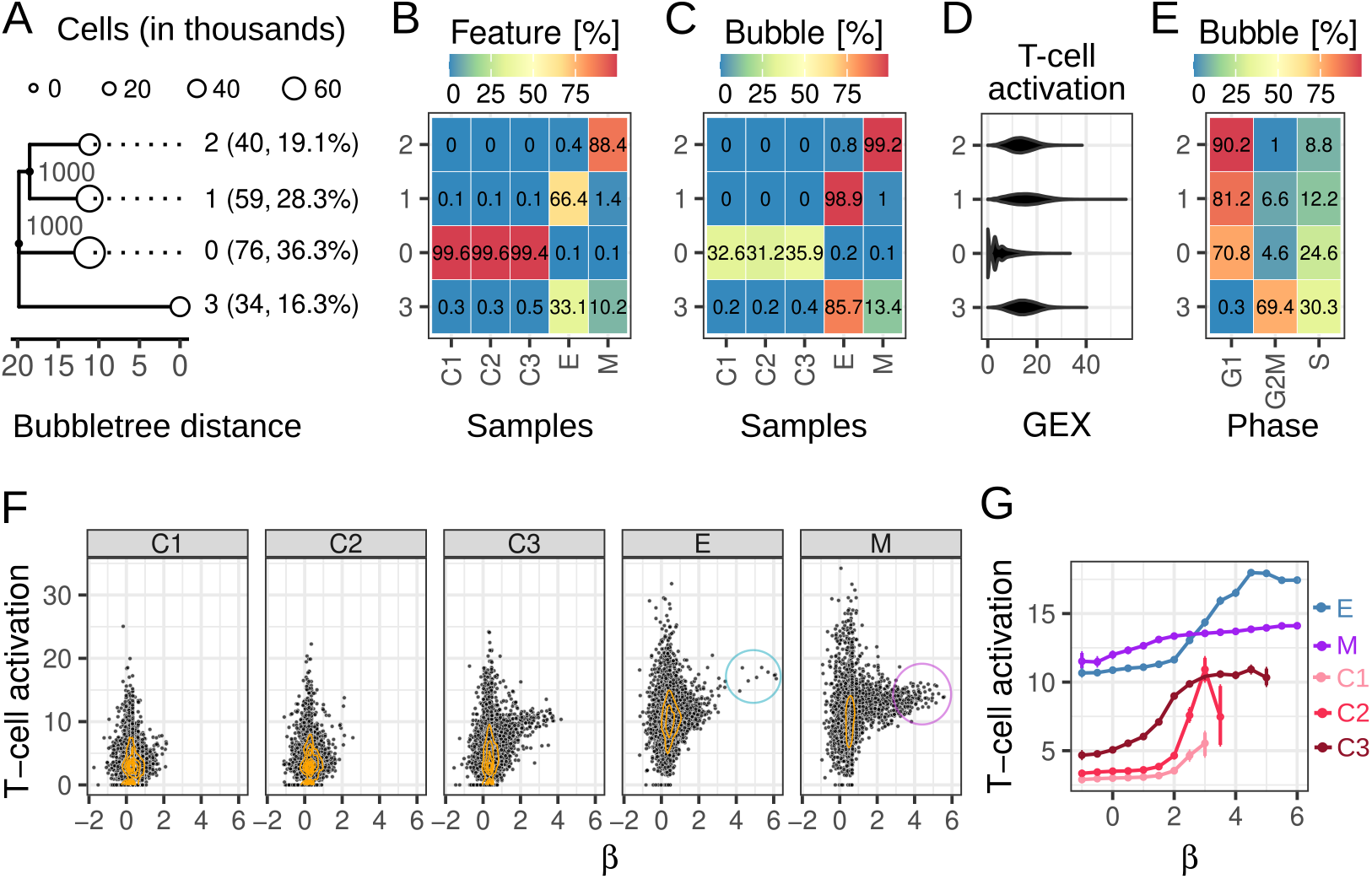
Integration of gene expression and TCR repertoires. (A) Annotated bubbletree (tree structure) of dataset 1. The bubbletree has five bubbles (white circles) as leaves. Bubble radii scale linearly with the number of cells in the bubbles. Bubble indices, absolute and relative cell frequencies are shown as labels between bubbletree and heatmap. Gray branch labels indicate perfect support of branches in 1,000 bootstrap iterations. (B) Heatmap columns are relative frequencies of cells from different samples, integrating to 100%. (C) Heatmap rows are within-bubble relative frequencies of different cell lines, integrating to 100%. (D) Distribution of normalized expression sum of all marker genes associated with T cell activation (x-axis) in cells from each bubble (y-axis). (E) Heatmap rows are within-bubble relative frequencies of cell cycle phases, integrating to 100%. (F) Mean T cell activation score (y-axis) versus *β* coefficient (x-axis) for Communities on the Joint graph (CJs) (dots) across five repertoires (panels). Orange contours show 2D density of CJs. (G) Mean T cell activation score (y-axis) for T cells in CJs grouped by *β* coefficients (x-axis; sliding window: midpoints *β* = 1 to 6 in 0.5 increments, window size 2). Error bars: 95% HDIs from 1,000 bootstraps. Colors: TCR repertoires.

Differential gene expression analysis revealed marked differences between stimulated (bubbles 1, 2, 3) and unstimulated (bubble 0) cells. Stimulated clusters showed upregulation of genes associated with T-cell activation (GO:0042110, Fig. 2D) and chemotaxis (GO:0010818). Importantly, on average we observed higher level of T cell activation in bubble 3 compared to bubbles 1 and 2. Meanwhile, bubble 0 showed upregulation of canonical naive T cell markers, including *CCR7, TCF7*, and *IL7R* (Supplementary Figure S5).

Bubbles were differentially enriched in cell cycle phases: bubbles 1 and 2 were predominantly in G1 phase (80% and 90% of cells, respectively), with the remainder in S or G2M phase (Fig. 2E). A cluster dominated by G1 suggests these T cells are quiescent, minimally activated, or resting T cells. On the other hand, bubble 3 was enriched in cells in G2M (70%) and S (30%) phases, characteristic of recently activated effector T cells undergoing clonal expansion (Fig. 2E), and consistent with the high T cell activation of cells from the stimulated repertoires (Fig. 2C-D). Bubble 0 cells were predominantly in G1 phase (71% of cells), with the remainder in S or G2M phase (Fig. 2E).

Notably, EBV stimulated CJs with high expansion (e.g., *β* > 3), including e1–e5, showed significantly elevated T cell activation scores compared to CJs with *β* ≤ 3 (Fig. 2F–G). Differential expression analysis confirmed the upregulation of many canonical activation and exhaustion markers (e.g., *NKG7, CCL5, GNLY, LAG3, TIGIT, IL2RA, CD70, CD2*) in *β* ≥ 3 CJs (Supplementary Figure S5). Conversely, while the MART1-stimulated repertoire M had elevated T cell activation levels relative to controls, the scores of its CJs increased only modestly with increasing *β*. In repertoire C3 activation increased with *β*; however, compared to repertoire E there were substantially fewer differentially expressed genes (DEGs) associated with T cell activation, with roughly balanced numbers of upregulated and downregulated markers between *β* > 3 and *β* ≤ 3 CJs (Supplementary Figures S6 and S7). This contrasts sharply with repertoire E, where *β* > 3 CJs showed pronounced upregulation of activation markers. Finally, repertoires C1 and C2 contained negligible numbers of *β* > 3 CJs (zero in C1, one in C2); however, in these repertoires, we still observed increased signatures of T cell activation associated with CJs having large *β* values.

### Tracking the dynamics of TCR CJs in lung cancer patients treated with CTLA-4 blockade and radiotherapy

To test whether *ClustIRR* can identify biologically meaningful expansion or contraction of TCR CJs in longitudinal TCR*β* repertoire data, we analyzed data from peripheral blood of 20 lung cancer patients treated with cytotoxic T lymphocyte antigen-4 (CTLA-4) checkpoint blockade and radiotherapy (RT) (***Formenti et al., 2018***). Between day 0 (before treatment) and day 22 (on treatment), *ClustIRR* identified a significantly greater fraction of expanding CJs (*β*_*day22*_> *β*_*day0*_ with non-overlapping 95% HDIs) in patients with complete or partial response (CR/PR) relative to those with stable or progressive disease (SD/PD) (overall difference = 0.075%, 95% HDI = [0.021, 0.146] (Fig. 3A, Supplementary Figure S9 and S10). These results suggest that early dynamic shifts in TCR CJs, particularly expansions, may serve as predictive biomarkers of treatment response in patients receiving CTLA-4 blockade and RT (***Formenti et al., 2018; Rudqvist et al., 2018***). Leave-one-out cross-validation (LOOCV) confirmed the robustness of this finding to the exclusion of individual samples (Supplementary Figure S11).

**Figure 3.**
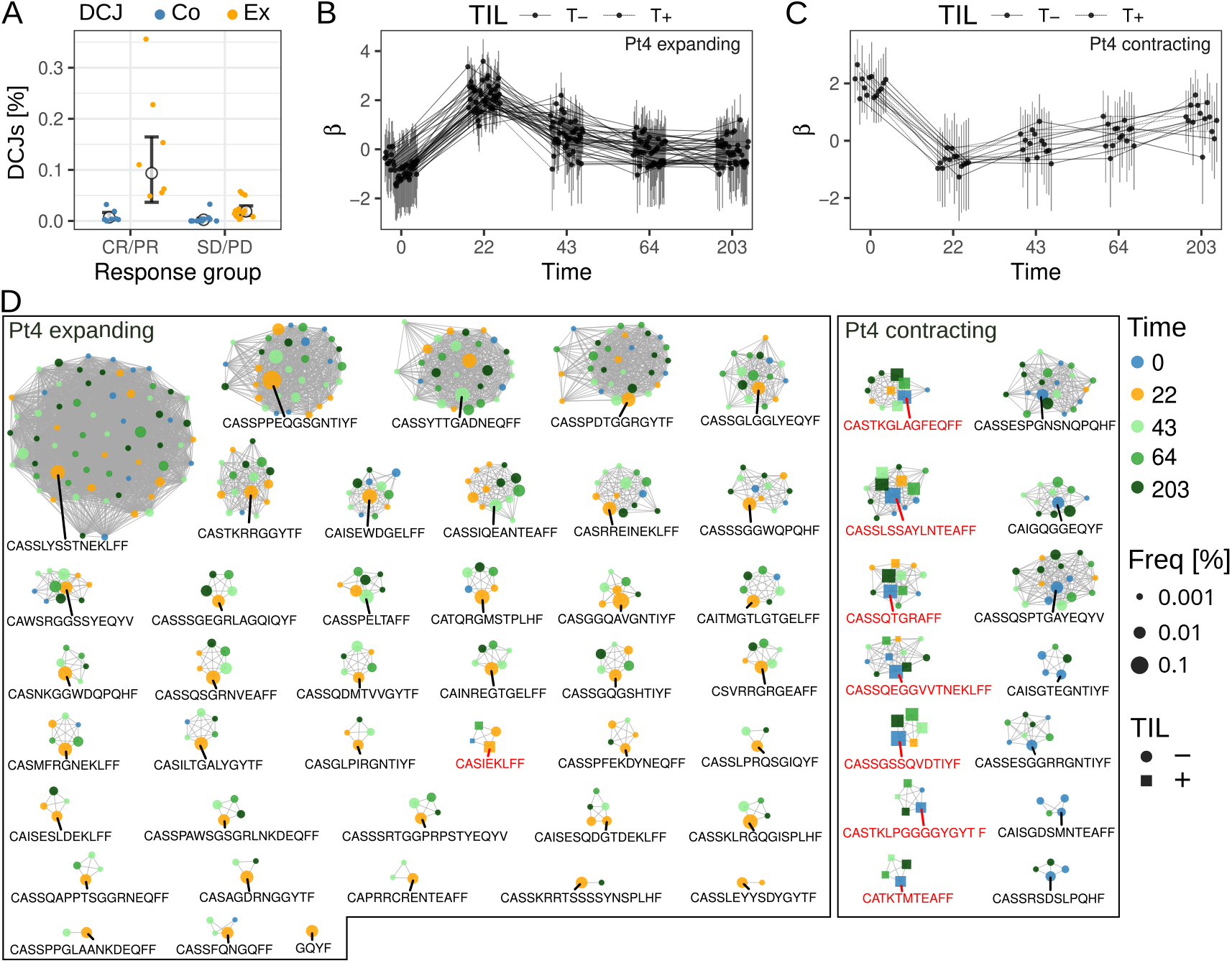
Analysis of longitudinal TCR data in lung cancer patients treated with CTLA-4 checkpoint inhibitors. (A) Changes of Differentially occupant CJs (DCJs) between day 0 and day 22. Percentage of expanding (Ex, blue dots) and contracting (Co, orange dots) CJs in responders (CR/PR) and non-responders (SD/PD), including the overall mean percentages (large black dot) and 95% HDIs (black error bars) in each group. (B/C) Mean CJ intensity (*β*, y-axis) and 95% HDI intervals (vertical error bars) for expanding (B) and contracting (C) DCJs in patient Pt4 across five timepoints (x-axis). Dashed and solid lines show CJs with (+) and without (-) tumor TCRs, respectively. (D) Network visualization of 41 expanding and 14 contracting DCJs in Pt4. Nodes represent clonotypes (colored by timepoint; sized by relative clonal expansion on logarithmic scale). Squares: tumor-infiltrating TCRs (T+); circles: non-tumor-infiltrating TCRs (T-). Red labels: CDR3*β* sequences of T+ clonotypes. The largest clonotype in each CJ is labeled with its CDR3*β* sequence. Edges connect clonotypes with similar CDR3*β*s.

*ClustIRR* further enables longitudinal tracking of TCR CJ expansion dynamics through timepoint resolved estimates of the CJ intensity parameter *β*, quantified by posterior means and 95% HDIs (points and vertical error bars in Fig 3B). In dataset 2, TCR*β* repertoires for 15 patients were sampled only at day 0 and day 22, while the remaining 5 patients had additional timepoints (Supplementary Figure S8). For example, in patient Pt4 (a treatment responder), TCR*β* repertoires were profiled at five timepoints: days 0, 22, 43, 65, and 203. By investigating the dynamics of expanding and contracting Differentially occupant CJs (DCJs), we identified two key patterns (Supplementary Figure S12). First, between days 0 and 22, DCJs occupancy changed markedly, with drastic shifts in *β* means and no overlap in their 95% HDIs. Second, between days 43 and 203, DCJs occupancies reverted to baseline (pre-treatment) levels in both expanding and contracting CJs (Fig. 3B, Supplementary Figure S12).

TCR*β* repertoires from two pre-treatment brain metastases in Pt4 enabled classification of peripheral blood TCRs as tumor-infiltrating (T+) if identical CDR3*β* sequences appeared in both compartments, or non-infiltrating (T-) when restricted to blood. Expanding or contracting DCJs containing T+ TCRs were subsequently categorized as T+ DCJs, while those lacking T+ TCRs were designated T-DCJs (dots connected by solid and dashed lines, respectively, in Fig. 3B). Notably, during days 0-22, 7 of 14 contracting DCJs were T+, potentially representing depletion of pre-existing tumor-antigen associated TCRs, while 1 of 41 expanding DCJs were T+, suggesting therapy-driven expansion of TCRs (Fig. 3C). These patterns are visually evident in network representations where T+ clonotypes localize to contracting CJs (Fig. 3C), though experimental validation of antigen specificity is needed for confirmation.

These results align with qualitative trends described in the original study (***Formenti et al., 2018***), but *ClustIRR* extends these findings by (i) quantifying uncertainty in longitudinal DCJ dynamics through Bayesian posterior estimates and (ii) broadening the analysis from individual TCR clonotypes to CJs of TCRs sharing similar CDR3*β* sequences, thereby capturing robust clonal dynamics.

### Quantifying TCR repertoire similarity and diversity across biological groups

Human and murine IRRs are dominated by private clonotypes but they should still be more similar within each species than between species. We hypothesized that *ClustIRR* with its community centric approach should reflect this similarity hierarchy better than a purely clonotype-centric approach.

To test this hypothesis, we generated synthetic CDR3*αβ* sequence pairs using OLGA (***Sethna et al., 2019***) and V(D)J generative models (***Marcou et al., 2018***) in humans and mice. This produced five human (H1-H5) and five mouse (M1-M5) TCR repertoires, each containing 10,000 clonotypes, with the five repertoires per species serving as biological replicates. We selected this multi-species strategy because systematic differences are known to exist between human and murine immune repertoires (***Madi et al., 2017***), owing to differential evolutionary challenges and the distinct sets of germline gene segments for V(D)J recombination.

The synthetic repertoires were very diverse, as expected from V(D)J recombination, with generation probabilities spanning tens of orders of magnitude (***Sethna et al., 2019***). No two clonotypes shared both CDR3*α* and CDR3*β* sequences, resulting in pairwise cosine similarity (CS) values of 0 (Figure 4A). While some clonotypes had identical CDR3*α* or CDR3*β* sequences, none shared both, highlighting the challenges faced by clonotype-centric approaches in analyzing multi-IRR data coherently. This problem is exacerbated by low sequencing depth in single-cell TCR sequencing.

**Figure 4.**
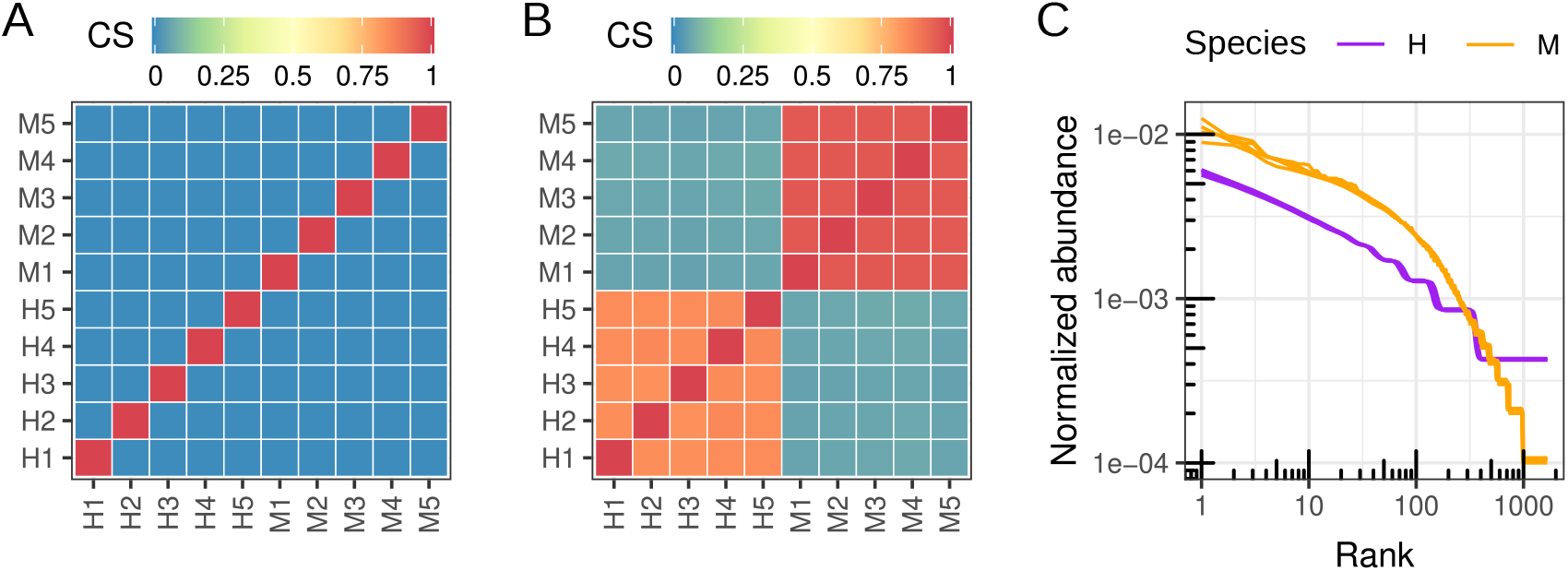
Analysis of TCR repertoires from humans and mice. (A) Heatmap with clonotype-centric cosine similarity (CS) between pairs of TCR repertoires. (B) Heatmap with community-centric CS between pairs of TCR repertoires. Color-coded tiles represent CS values. (C) Log-log plot of normalized rank abundance distributions (NRADs) of TCR repertoires from humans (H, purple) and mouse (M, orange) repertoires. Each line represents an individual repertoire.

With its community-centric design, *ClustIRR* should cope better with this data. *ClustIRR* first con-structed a joint graph of 100,000 nodes (clonotypes) connected by ≈10 million edges, with weights representing CDR3*αβ* sequence similarity (Methods). Leiden clustering (Potts model; resolution *r* = 1) identified 14,160 CJs. CJ distribution analysis revealed 5,742 (≈40%) private CJs exclusive to individual repertoires and 3,663 CJs (≈26%) shared between two repertoires. As expected, 991 CJs (≈7%) were shared across all five human repertoires, and 906 CJs (≈6.5%) were shared across all five mouse repertoires. However, only 47 CJs (≈0.3%) were shared across all ten repertoires from both species, underscoring the rarity of cross-species convergence.

To quantify repertoire similarity, we calculated pairwise CS between all ten TCR repertoires based on their CJ distributions. The results revealed stark differences between intra- and interspecies similarity. Human repertoires showed high intra-species similarity, with CS values ranging from 0.83 to 0.85 (Figure 4B), while mouse repertoires were even more similar to each other (CS: 0.94–0.95). In contrast, inter-species similarities were very low with CS values of 0.06–0.07. These findings are consistent with known evolutionary and immunological differences between human and murine IRRs, and they validate the biological relevance of *ClustIRR*’s community-centric approach.

We further assessed TCR repertoire diversity using normalized rank abundance distributions (NRADs) (***Saeedghalati et al., 2017***), which capture both the dominance structure (whether a few CJs overwhelmingly dominate the repertoire) and the overall diversity (how evenly CJs are distributed across the population). Mouse repertoires had a “top-heavy” distribution, with the top 100 CJs accounting for disproportionately high normalized abundance, while human repertoires showed flatter NRADs, indicating greater diversity (Figure 4C). Within each species, NRADs were almost identical between biological replicates.

Thus, the community-centric framework of *ClustIRR* makes similarities between different but comparable repertoires accessible, even if all observed clonotypes are private, be it because of biological combinatorics or because of limited sequencing depth.

### Convergent repertoire differences between species

To identify convergent repertoire differences between humans and mice, we applied *ClustIRR*’s hierarchical Bayesian model for differential CJ occupancy (Methods: model ***M***_*h*_). Building on the repertoire-specific effects on CJ occupancy described by parameters *β*_*i*_, this hierarchical framework additionally estimates overall species-specific CJ occupancy parameters (*µ*_*j*_; dots in Figure 5A), where *j* indexes biological conditions (human or mouse). By modeling abundance at both the individual and group levels, this approach distinguishes conserved species-level trends from individual repertoire variability, enabling the robust detection of systematic shifts in CJ representation between species.

**Figure 5.**
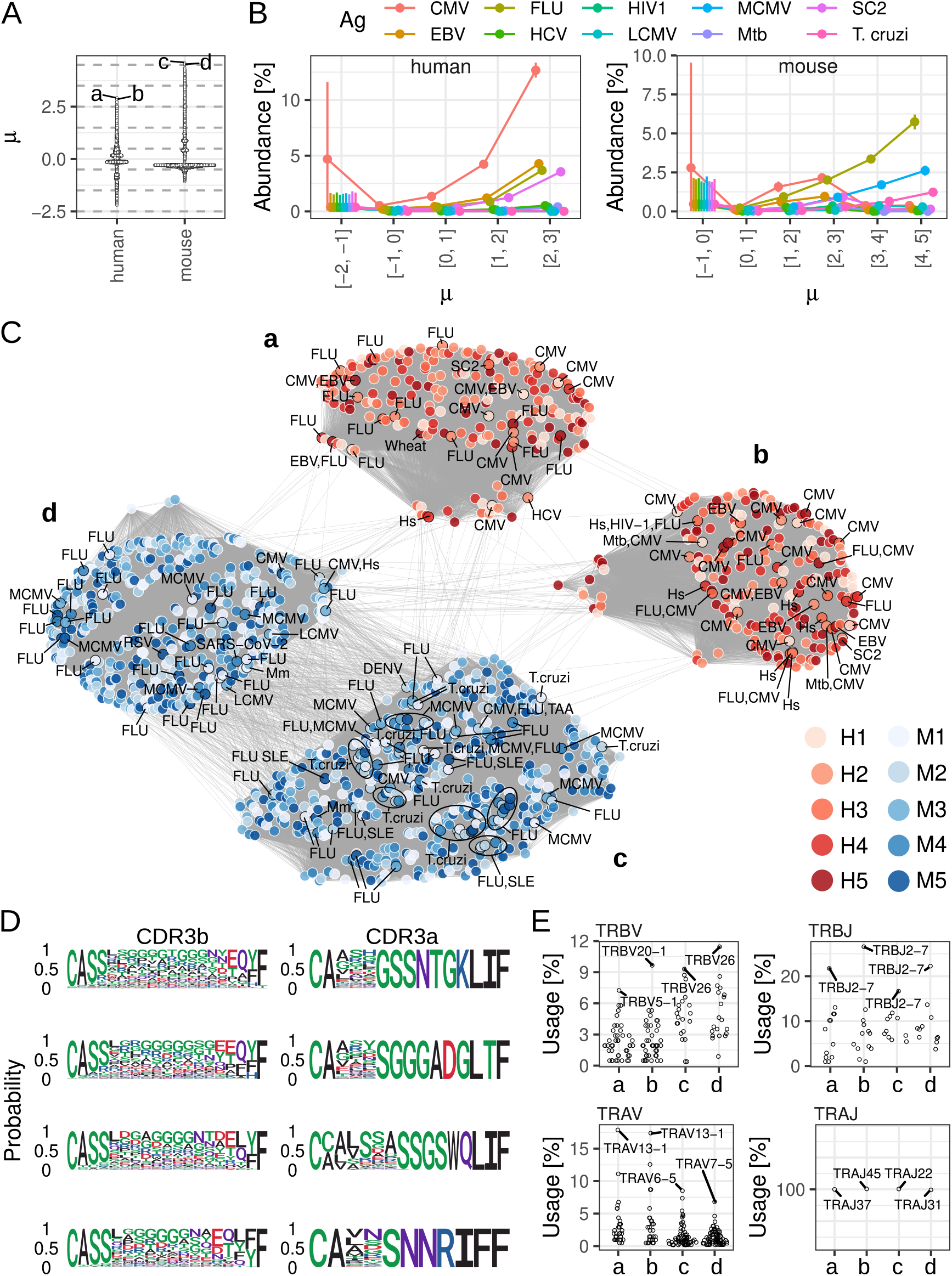
Differentially occupant CJs (DCJs) between humans and mice. (A) Relative occupancy of CJs (dots) in humans and mice, represented by the mean of *µ* (y-axis). Horizontal dashed lines split the CJs into six groups used in panel B. Two most expanded in humans and mice are labeled (a–d). (B) Mean percentage of clonotypes recognizing specific antigens (color-coded) within CJ groups (x-axis) and the corresponding 95% HDIs (vertical error bars). The groups are ordered by increasing *µ*. Groups with no antigen-specific clonotypes were omitted. (C) Network of four DCJs, shared across all repertoires of one species (a–b in humans, c–d in mice) but not the other. Nodes represent TCR clonotypes from humans (reds) and mice (blues). Gray edges connect clonotypes sharing similar CDR3 sequences. Antigen-specific clonotypes are shown as nodes with black borders and labeled with the names of the antigen species. (D) Sequence logos of more conserved CDR3*α* motifs and more divergent CDR3*β*s within each CJ (rows: CJs a–d) (E) Relative usage (y-axis) of TRBV/TRBJ/TRAV/TRAJ gene segments (panels) in each CJ (x-axis). The most frequently used gene segments are labeled.

The distribution of *µ*_*j*_ revealed pronounced shoulders toward large values in both species, indicating the presence of large, highly occupant CJs enriched in clonotypes from each species. To investigate whether these CJs are functionally relevant, we binned CJs into seven groups based on their *µ*_*j*_ means (horizontal bins in Figure 5A). For each bin, we computed the probability that clonotypes within the CJ recognize antigens from ten pathogenic species with abundant annotations in VDJdb (Figure 5B). We observed a clear enrichment pattern: CJs with higher *µ*_*j*_ values in humans were increasingly enriched for clonotypes targeting human-specific pathogens, including CMV, EBV, Influenza A, and SARS-CoV-2. Similarly, in mice, high-*µ*_*j*_ CJs were enriched for clonotypes recognizing mouse-specific pathogens such as FLU, MCMV, and *Trypanosoma cruzi*.

Because these repertoires are synthetic—generated using models trained on non-selected TCRs to capture intrinsic V(D)J recombination biases—this enrichment cannot be attributed to immune selection or antigenic pressure. Instead, it reflects the convergent generation of functionally relevant CDR3 motifs driven by intrinsic biases in V(D)J recombination. To quantify this, we investigated the relationship between CJ size and the mean probability of generation (***P**_gen_*) of their constituent clonotypes, computed by OLGA (***Sethna et al., 2019***). As expected, we observed a positive correlation: large CJs had high average ***P***_*gen*_ (Supplementary Figure S13A). This indicates that CDR3 sequences with high ***P***_*gen*_ are statistically more likely to be generated, leading to the convergence of similar sequences into “public” CJs.

To validate that this generation bias underpins the observed antigen enrichment, we compared the ***P***_*gen*_ distributions of CDR3 sequences specific for common pathogens against non-specific CDR3s (according to VDJdb annotations). In both humans and mice, CDR3*α* sequences specific for common antigens had significantly higher average ***P***_*gen*_ compared to non-specific CDR3*α*s (Supplementary Figure S14). In contrast, this pattern was not observed for CDR3*β*; antigen-specific and unspecific CDR3*β*s showed overlapping ***P***_*gen*_ distributions, with significantly lower values overall compared to CDR3*α*, consistent with the greater diversity of the *β* chain (Supplementary Figure S14). These results suggest that intrinsic generation biases in the *α* chain are a primary driver of the high occupancy, antigen-enriched CJs.

We further analyzed the network structure and sequence features of the two most occupant CJs in humans (a, b) and mice (c, d) (Figure 5C). While these CJs were shared within species, they were largely non-overlapping across species. Sequence logos (Figure 5D) emphasized conserved CDR3*α* motifs, particularly at the 3’ end where joining (J) gene segments are incorporated, whereas CDR3*β* regions were more variable. Gene usage analysis (Figure 5E) confirmed this structural asymmetry: while each CJ used a mixture of TRBV, TRBJ, or TRAV genes was observed, it used only one TRAJ gene segment. This suggests that the conserved germline-encoded core of CDR3*α* sequences, dictated by TRAJ selection, is a key determinant of CJ identity.

This pattern generalized to all CJs within the highest *µ*_*j*_ bins (131 CJs in humans with *µ*_*j*_ ≥ 2 and 35 CJs in mice with *µ*_*j*_ ≥ 4; Supplementary Figure S15). Importantly, we observed completely disjoint posteriors for *µ*_*j*=human_ versus *µ*_*j*=mouse_ for most of these CJs, indicating clear differential CJ occupancy. Specifically, mean *µ*_*j*=human_> *µ*_*j*=mouse_ in 113 out of 131 CJs (enriched in humans relative to mice) and *µ*_*j*=mouse_> *µ*_*j*=human_ in all 35 CJs (enriched in mice relative to humans). Nearly all of these CJs had conserved CDR3*α* motifs paired with diverse CDR3*β*s. Crucially, although the CJs used different TRAJ gene segments, all were enriched for multiple human-specific or murine-specific pathogens, respectively.

We next asked whether this germline-driven convergence extends to the *β* chain across the broader set of DCJs. Among 1,000 CJs with completely disjoint posteriors for *µ*_*j*=human_ versus *µ*_*j*=mouse_ (including 114 enriched in humans and 886 in mice), conserved CDR3*β* motifs were rare. Only 173 CJs (all except for one having higher occupancy in mice) had conserved CDR3*β* motifs with *µ*_*j*=mouse_ ∈ [1.7, 4.3]. This scarcity of convergent *β* CJs likely arises from the generally lower generation probabilities (***P***_gen_) of CDR3*β* sequences in both humans and mice. Importantly, in these CJs we observed the same pattern: conserved TRBJ gene segments were key determinants of CJ identity (Supplementary Figure S16). However, even in these rare *β*-driven CJs, we did not observe significant enrichment for murine-specific pathogens, consistent with the overlapping ***P***_gen_ distributions for antigen-specific and unannotated CDR3*β* sequences (Supplementary Figure S14). Thus, while the *α* chain drives functional convergence through germline bias, the *β* chain remains highly diverse and less functionally constrained in the absence of selection.

Together, these findings reveal a fundamental asymmetry where germline biases preferentially converge antigen-specific CDR3*α* sequences while maintaining CDR3*β* diversity, which *ClustIRR* quantitatively identified through specific CJ examples, enabling mechanistic interpretation. With its hierarchical model that distinguishes individual variability from group level trends with probabilistic quantification of CJ occupancy and their uncertainty, *ClustIRR* provides a robust framework for uncovering such systematic biases in diverse case studies beyond species comparisons (e.g., treated vs. untreated individuals, disease vs. healthy cohorts).

### Benchmarking

To evaluate the computational scalability and parameter sensitivity of *ClustIRR*, we performed bench-marking analyses varying the repertoire depth and the global minimum identity (GMI) threshold using paired TCR *αβ* CDR3 sequences. GMI specifies the minimum pairwise amino acid sequence identity required between two CDR3 sequences to retain an edge in the similarity graph. We measured the CPU time for the primary processing function (`clustirr`, dominated by similarity search) and the CJ detection function (`detect`_`communities`), peak random access memory (RAM) usage, and the resulting number of detected CJs.

First, we assessed the impact of repertoire depth using subsampled datasets ranging from 1,000 to 100,000 paired clonotypes, fixed at a GMI threshold of 0.8 (Supplementary Figure S15A– B). As expected, computational time scaled super-linearly with input size, reflecting the pairwise nature of sequence similarity calculations. The primary processing time increased from approximately 17 seconds for 1,000 clonotypes to 2.4 hours for 100,000 clonotypes (Supplementary Figure S15A). CJ detection time also increased with depth, from 1.7 seconds to 11 minutes, but remained a minor fraction of the total runtime. Peak memory usage scaled similarly, with clustirr requiring approximately 0.1 GB for 1,000 clonotypes and 13 GB for 100,000 clonotypes, while `detect_communities` required 0.03 GB and 7 GB, respectively (Supplementary Figure S15B). Notably, the number of detected CJs grew linearly with clonotypes on the log-log scale in both CPU and RAM panels, indicative of power-law scaling. This suggests that CJ discovery increases predictably with sampling effort rather than saturating, reflecting the underlying hierarchical structure of the repertoire where new CJs continue to emerge at higher sequencing depths.

Second, we evaluated the effect of the GMI threshold (0.6 to 0.95) on a fixed repertoire of 5,000 paired clonotypes (Supplementary Figure S15C). The primary processing time remained stable across GMI values (170 seconds), confirming that the computational bottleneck lies in the initial similarity search, which precedes threshold filtering. In contrast, CJ detection time decreased markedly with increasing GMI, dropping from 18 seconds at GMI=0.6 to 0.7 seconds at GMI=0.95. This reduction corresponds to a sparser similarity graph at stricter identity thresholds. Concurrently, the number of detected CJs increased eight-fold (from ≈590 to ≈4,950) as the GMI threshold rose, reflecting increased fragmentation of the graph into smaller, tighter sequence groups. These results demonstrate that while repertoire depth dictates overall runtime and memory, the GMI parameter allows users to tune CJ resolution and downstream clustering speed without affecting initial processing costs.

## Discussion

We have shown above that *ClustIRR* is a capable computational framework for uncertainty-aware quantitative analysis of TCR CJ occupancy. It provides a unique approach to IRR analysis by combining graph-based community detection with hierarchical Bayesian modeling, *ClustIRR* addresses fundamental challenges in repertoire analysis: clonotype sparsity, biological noise, and the need for rigorous uncertainty-aware quantification. Our results demonstrate that *ClustIRR* successfully identifies antigen-specific TCR CJs, links clonal expansion to transcriptional activation states, tracks longitudinal dynamics in cancer immunotherapy, and reveals intrinsic evolutionary biases in TCR generation that pre-adapt repertoires for pathogen recognition.

Traditional clonotype-centric approaches struggle with the extreme diversity of TCR repertoires, where identical sequences are rarely shared across samples. By aggregating clonotypes into sequence similarity CJs, *ClustIRR* enables meaningful comparisons even if no clonotypes are shared between samples. Our synthetic repertoire analysis demonstrated this clearly: while clonotype-level similarity between human and mouse repertoires was zero, community-level similarity revealed substantial intra-species conservation and distinct inter-species architecture. This community-centric approach aligns with evidence that TCRs with similar sequences often recognize the same epitopes (***Glanville et al., 2017; Dash et al., 2017***), an insight that *ClustIRR* exploits to provide a robust framework for detecting convergent immune responses.

Furthermore, *ClustIRR*’s hierarchical Bayesian Dirichlet-Multinomial model explicitly accounts for overdispersion, generating posterior distributions rather than point estimates. This enables researchers to distinguish true biological expansion from stochastic fluctuations–critical for interpreting repertoire dynamics in settings with limited sampling depth, as shown in our longitudinal cancer immunotherapy analysis where early CJ expansions predicted treatment response.

In practical applications, *ClustIRR* proved effective across diverse biological contexts. In single-cell datasets, we identified antigen-specific CJs expanded upon stimulation, and we could link these expansions to transcriptional activation signatures. This integration of transcriptomes and TCRs is a key advantage that enables functional interpretation of expanding CJs by characterizing the activation state of the underlying cells.

In longitudinal bulk sequencing data, *ClustIRR* detected early dynamic shifts in CJ occupancy that could serve as predictive biomarkers of treatment response.

In the third application, we used *ClustIRR* to analyze simulated repertoires from human and mouse, and we could demonstrate the framework’s ability to detect inter-species shifts in TCR repertoires, namely species-specific CJ structures enriched for pathogen-specific motifs even without antigenic selection in individual organisms. As a likely mechanism behind these shifts could be species-intrinsic V(D)J recombination biases. This finding suggests that generation probabilities (***P***_gen_) from models like OLGA could serve as informative priors in Bayesian repertoire analysis, particularly for *α*-chains where germline encoding is strongest.

Importantly, the framework’s utility extends to complex clinical cohorts beyond the benchmark applications presented here. In a companion study (***Kitanovski et al., 2026***), we used *ClustIRR* to analyze paired primary melanoma tumors and sentinel lymph nodes from 24 treatment-naive patients. This investigation revealed profound spatial compartmentalization of TCR repertoires, where primary tumors exhibited reduced diversity and pronounced clonal dominance compared to matched lymph nodes. Despite the high degree of clonotype privacy between compartments and individuals, *ClustIRR*’s community-centric approach successfully identified recurrent TCR communities enriched for shared melanoma antigens (e.g., MART-1) as well as patient-specific neoantigen signatures. This real-world application demonstrates *ClustIRR*’s robustness in extracting biologically meaningful signals from sparse, high-dimensional clinical data where traditional clonotype-centric methods often fail to resolve shared immune architectures.

While *ClustIRR* offers a ready-to-use pipeline for sequence clustering and statistical analysis, its modular design enables interoperability with other established tools. Users can import CJ definitions from methods such as TCRdist3/TCRdist (***Dash et al., 2017; Mayer-Blackwell et al., 2021***), GIANA (***Zhang et al., 2021***), GLIPH/GLIPH2 (***Glanville et al., 2017; Huang et al., 2020***), iSMART (***Zhang et al., 2020***), TCRclust (***Valkiers et al., 2021***), or deepTCR (***Sidhom et al., 2021***) for downstream Bayesian modeling within *ClustIRR*, facilitating method comparison and leveraging specialized clustering algorithms. This flexibility is particularly valuable given several implementation considerations.

First, while building a joint graph across different repertoires and conditions is a key advantage of *ClustIRR* and can be performed efficiently with our BLAST-based similarity strategy, computational challenges arise when analyzing hundreds of repertoires with hundreds of thousands of clonotypes each (e.g., the Emerson dataset (***Emerson et al., 2017***)). In such scenarios, the pairwise similarity calculations and graph construction become computationally demanding. This limitation can be mitigated by applying multicore execution, which is already implemented in *ClustIRR*, or by utilizing the aforementioned modular design to integrate more efficient clustering approaches such as GIANA (***Zhang et al., 2021***) or ClustTCR (***Mayer-Blackwell et al., 2021***) for the initial CJ detection step.

Second, our hierarchical Bayesian model is parameter-rich, which ensures flexibility but comes at a computational cost. When analyzing many CJs, repertoires, and biological conditions simultaneously, substantial RAM and CPU time are required. For the datasets presented here, runtime ranged from hours to one day. However, we can envision scenarios with larger cohort sizes or deeper sequencing where computational demands could become prohibitive. Fortunately, emerging techniques such as amortized Bayesian inference (***Li et al., 2024***) and variational inference (***Blei et al., 2017***) could in the future substantially reduce computational burden while maintaining uncertainty quantification.

Third, *ClustIRR*’s built-in CJ detection depends on similarity thresholds that may require tuning for specific applications. Finally, while we demonstrated *ClustIRR* on TCR repertoires, application to BCR repertoires requires validation given differences due to somatic hypermutation.

In conclusion, *ClustIRR* provides a rigorous, uncertainty-aware framework for quantitative TCR repertoire analysis. By moving beyond clonotype-centric approaches to community-level inference, *ClustIRR* enables detection of convergent immune responses, integration with functional genomics data, and discovery of fundamental principles of repertoire architecture. Its modular design ensures compatibility with existing workflows, allowing researchers to combine specialized clustering solutions with robust statistical modeling. As TCR sequencing becomes increasingly routine in clinical and research settings, tools like *ClustIRR* will be essential for translating repertoire data into biological insight and therapeutic opportunity. *ClustIRR* is available as an open-source R package via Bioconductor.

## Methods and Materials

We present a computational framework that integrates high-throughput TCR sequencing data with Bayesian probabilistic modeling to quantify differences in the structure and dynamics of TCR repertoires under uncertainty. This section is organized to first describe the datasets and then detail the *ClustIRR* framework, which combines graph-based community detection with hierarchical Bayesian modeling. The overall analytical workflow is summarized in Figure 6.

**Figure 6.**
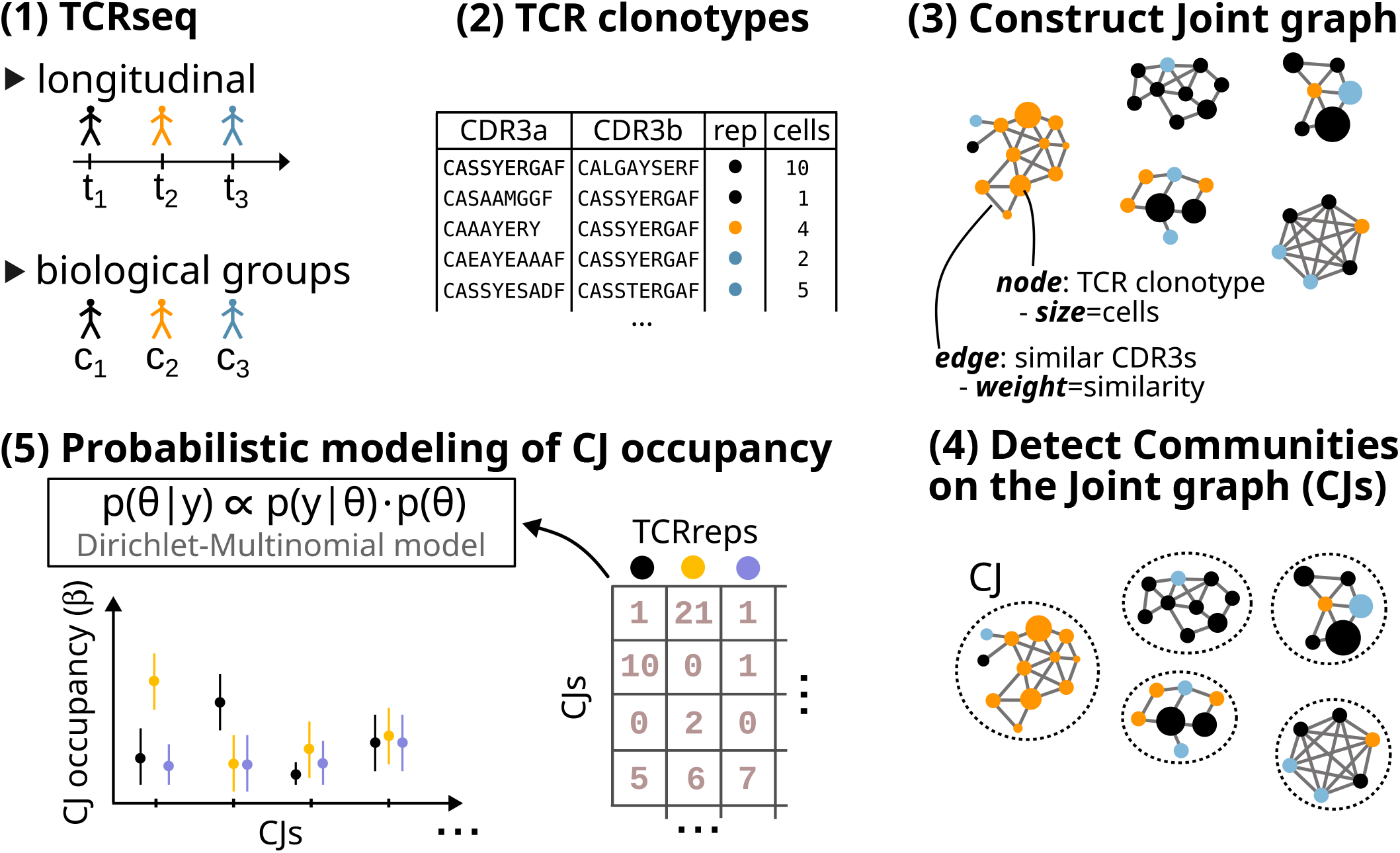
ClustIRR workflow. (1) TCR sequencing (TCRseq) data are collected from multiple repertoires across longitudinal time points or biological conditions. 2) The main *ClustIRR* input is a table of TCR clonotypes for each repertoire, defined by CDR3*α* and CDR3*β* amino acid sequences and clonal sizes (cell counts). (3) *ClustIRR* constructs a joint similarity graph where nodes represent clonotypes and weighted edges connect clonotypes based on pairwise CDR3*α* and CDR3*β* sequence similarity. This graph integrates clonotypes from all repertoires into a single structure. (4) Graph-based clustering algorithms (e.g., Leiden or Louvain) identify densely connected groups of clonotypes, defined as *Communities on the Joint graph* (CJs). The number of cells in each CJ is quantified to form a CJ occupancy matrix ^*k*×*r*^, where *k* indexes CJs and *r* indexes repertoires. This matrix is used to fit a hierarchical Bayesian Dirichlet-Multinomial model. The model estimates posterior distributions for relative CJ occupancies (*β* coefficients), providing mean estimates along with 95% credible intervals (CIs).

### Datasets

In this work, *ClustIRR* was evaluated with three datasets.

#### Dataset 1: single cell TCR*αβ* repertories of antigen-reactive human T cells

Dataset 1 contains single cell TCR*αβ* (scTCR) repertories obtained from Parse Biosciences (https://www.parsebiosciences.com/datasets/tcr-sequencing-of-1-million-antigen-reactive-human-t-cells-in-a-single-experime accessed on 17.10.2024). A detailed description of the data generation and processing protocol is provided by Parse Biosciences, and a summary is given in the following. The dataset contains TCR sequences of up to 1 million T cells isolated from 15 human donors’ PBMCs. The T cells were purchased from a vendor (https://www.criver.com/products-services/cell-sourcing/human-peripheral-blood/antigen-specific-t-cells?region=3601), and were stimulated with tumor, viral, and autoantigens. In addition, the dataset contains three TCR repertoires from unstimulated, healthy donors as controls. The samples were analyzed in a single experiment with the Evercode TCR Mega kit, and TCR specific libraries were sequenced on Illumina Novaseq X, followed by data processing with Parse Biosciences Analysis Pipeline v1.3.0.

The scTCR dataset contained data from 1,307,803 cells from 15 TCR repertoires (Supplementary Table 1). First, we removed 98,329 cells classified as multiplets. Furthermore, we removed 217,285 cells with missing TRAV or TRBV gene annotation, and 17 cells with non-productive TCRs. Each of the 992,170 remaining T cells had paired TCR*α* and TCR*β* chains, including their CDR3*α* and CDR3*β* amino acid sequences, and TRAV/TRAJ and TRBV/TRBJ genes. These cells were distributed across 100,980 clonotypes. For our analysis, we selected the 5 largest TCR repertoires in terms of T cell clonotypes, including: 3 unstimulated controls (C1, C2, and C3), and two repertoires stimulated with Epstein-Barr virus (EBV) and Melanoma Antigen Recognized by T cells 1 (MART1) antigens (TCR repertoires E and M, respectively).

Alongside the single cell TCRs, RNA-seq data was available, which was processed with the R-package *Seurat* (version 5.2.1). Quality control involved removal of cells with high relative frequency (>15%) of mitochondrial RNA, and more than 20,000 detected RNA transcripts or more than 5,000 detected genes per cell. Data processing involved normalization (function *LogNormalization* with scale.factor = 10,000), identification of 3,000 variable features (function *FindVariableFeatures* with selection.method = vst), data scaling while regressing out mitochondrial RNA content (function *ScaleData*), principal component analysis (PCA, with function *RunPCA*) and cell cycle phase prediction (function *CellCycleScoring*).

#### Dataset 2: TCR*β* repertoires from lung cancer patient PBMCs

Dataset 2 contains bulk TCR*β* repertoires from 20 lung cancer patients treated with CTLA-4 blockade (anti-CTLA-4 antibody, ipilimumab) and radiotherapy (RT) (***Formenti et al., 2018***). Treatment response was observed in each patient: two with complete response (CR), 5 with partial response (PR), 5 with stable disease (SD), and 8 with progressive disease (PD) (Supplementary Figure 5). TCR*β* sequencing was performed on the peripheral blood samples before treatment (day 0) and at one or multiple timepoints during treatment (Supplementary Figure 5). For patients Pt4, Pt32, Pt36 and Pt38, TCR*β* sequencing was performed on pretreatment (archival) tumor tissue. A detailed description of the data generation and processing protocol is provided in the original publication (***Formenti et al., 2018***), and a summary is provided in the following. DNA was isolated from PBMCs, and sequencing of TCR*β* CDR3 regions was performed using the ImmunoSEQ platform from Adaptive Biotechnologies, which measures the absolute cellular abundance of T cell clonotypes in each repertoire. The TCR sequence data were obtained from the ImmuneACCESS project repository of the Adaptive Biotechnology database (https://doi.org/10.21417/B7BW6X; downloaded in December 2024)

#### Dataset 3: simulated single cell TCR*αβ* repertories from humans and mice

Dataset 3 comprises ten TCR*αβ* repertoires from: (i) five simulated human repertoires (H1-H5), and five simulated mouse repertoires (M1-M5). The repertoires were generated using the OLGA algorithm (v1.2.4, function *olga-generate_sequences*) (***Sethna et al., 2019***). OLGA implements models of V(D)J recombination in healthy humans and mice, enabling simulation of TCR repertoires with CDR3*α* or CDR3*β* sequences. We created five TCR repertoires of humans (H1-H5) and five TCR repertoires of mice (M1-M5) by simulating 10,000 CDR3*α* and 10,000 CDR3*β* amino acid sequences in each repertoire. As recent analyses suggests that TCR chain pairing is nearly independent (***Dupic et al., 2019***), we joined the individual CDR3*α* and CDR3*β* sequences to generate CDR3*αβ* pairs. The five TCR repertoires per species were treated as biological replicates. Finally, we used OLGA’s function *olga-compute_pgen* to compute the generation probability (***P***_*gen*_) of each simulated CDR3 sequence, restricted to the corresponding V and J genes, according to the same generative V(D)J models used to generate the data.

### Analysis of TCR repertoires with *ClustIRR*

TCR repertoire analysis was performed with *ClustIRR* (R-package, version 1.9.38). The main input to *ClustIRR* is a table with TCR clonotypes from a TCR repertoire, where each clonotype contains three elements: CDR3*α* and CDR3*β* amino acid sequences and cell count (clonotype size). *ClustIRR* can also process inputs containing clonotypes with only CDR3*α* or CDR3*β* sequences and their clonal sizes to facilitate analysis of TCR repertoires generated by bulk TCR sequencing (e.g., Dataset 2). *ClustIRR* performs the following steps:

1. Compute similarities between T cell clonotypes
2. Construct a joint similarity graph (*J*)
3. Detect Communities on the Joint graph (CJs)
4. Analyze Differential CJ Occupancy
5. Between individual TCR repertoires with model M
6. Between groups of TCR repertoires from biological conditions with model M_h_
7. Inspect communities of interest

### Computing similarities between TCR clonotypes

*ClustIRR* aims to quantify the similarity between pairs of T cell clonotypes based on their CDR3 sequences. Given a TCR repertoire with *n* clonotypes, we have to perform *n*^2^ pairwise comparisons of CDR3*α* and CDR3*β* sequences. HT-seq technology allows us to profile TCR repertoires with over a million clonotypes (*n* > 10^6^) in a single study. Thus, pairwise CDR3 sequence comparisons using algorithms such as the Needleman-Wunsch global sequence alignment algorithm may require prohibitive computational resources.

To address this, *ClustIRR* employs Basic Local-Alignment Search Tool (BLAST, ***Altschul et al. (1990)***). Briefly, a protein database is constructed (method *makeblastdb*) from all CDR3 sequences, and each CDR3 sequence is used as a query (method *blastp*). This enables fast sequence similarity searches. By default, only CDR3 sequences matches with ≥ 80% sequence identity to the query are retained. This step reduces the computational and memory requirements, without impacting downstream CJ analyses, as CDR3 sequences with lower typically yield low similarity scores.

For matched CDR3 pair, an alignment score (*ω* ∈ ℤ) is computed using BLOSUM62 substitution scores (***Henikoff and Henikoff, 1992***) with gap opening penalty of −12 and gap extension penalty of −2. Identical or highly similar CDR3 sequence pairs receive large positive *ω* scores, while dissimilar pairs receive low or negative *ω*.

To normalize *ω* for alignment length, *ClustIRR* computes 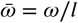, where is the alignment length yielding normalized alignment score 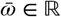 This normalization, also used in iSMART (***Zhang et al., 2020***), ensures comparability across CDR3 pairs of varying lengths.

*ClustIRR* also computes alignment scores for the CDR3 *core* regions (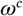 and 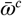). The CDR3 core, representing the central loop region with high antigen-contact probability (***Glanville et al., 2017***), is generated by trimming three residues from each end of the CDR3 sequence. Comparing 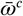 and 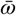 allows assessment of whether sequence similarity is concentrated in the core or flanking regions.

### Constructing a joint similarity graph (*J*)

*ClustIRR* uses the R-package *igraph* (version 2.1.1) to create graphs for each TCR repertoires. T cell clonotypes are represented as graph nodes with undirected edges connecting pairs of nodes based on CDR3*α* and CDR3*β* similarity scores (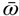 or 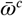). Additionally, *ClustIRR* computes similarities between CDR3 sequences of clonotypes from different repertoires (as in Section “Computing similarities between TCR clonotypes”), creating new edges between nodes from different graphs with 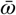 and 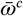 as edge attributes. The joint graph *J* is structured as an igraph object.

To prioritize memory efficiency while also improving computational time with large datasets, *ClustIRR* includes an optional *k*-nearest neighbor graph (*k*-NNG) algorithm. When enabled, connections are restricted to a maximum of *k* highest-scoring neighbors per chain (CDR3*α* or CDR3*β*) based on similarity metrics (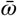 or 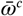), reducing edge count and memory usage. In this mode, when integrating a new repertoire into *J*, existing nodes may form edges to up to 2*k* new neighbors (*k* for *α* and *k* for *β* chains), preserving high-similarity connections both within and between repertoires while maintaining scalability. Using the default setting (*k* = 50), this approach yields downstream results comparable to the full graph but reduces computational resources, as benchmarked on repertoires of 5,000–100,000 clonotypes (Supplementary Figure X). For larger repertoires, we recommend increasing *k* to ensure concordance with the full graph. Accordingly, we employed a full graph analysis for Datasets 1 and 2, and the *k*-NNG approach (*k* = 50) for Dataset 3.

### Graph-based CJ detection

*ClustIRR* identifies CJs in *J* using graph-based community detection (GCD) algorithms such as Leiden (***Traag et al., 2019***), Louvain (***Blondel et al., 2008***) or Infomax (***Rosvall and Bergstrom, 2008***). First, the similarity score between T cell clonotypes *i* and *j* is defined as the average CDR3*α* and CDR3*β* alignment scores:

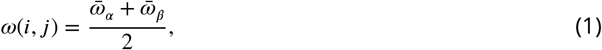

where 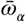 and 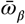 are the alignment scores for the CDR3*α* and CDR3*β*, respectively. If a chain is missing, its alignment score is set to 0.

For Datasets 1-3, Leiden clustering was applied on the weighted graph as implemented in the R-package *igraph* (version 2.1.1), with a Constant Potts Model (CPM) quality function (***Traag et al., 2011***), resolution parameter set to 1, and 1,000 optimization iterations, generating a CJ occupancy matrix *y*^*k*×*r*^ with entries representing the number of T cells in *k* (for *k* = 1, …, *K*) and repertoire *r* (for *r* = 1, …, *R*). In designs with multiple repertoires per biological condition (e.g., Dataset 3), repertoires are grouped by condition *b* (for *b* = 1, …, *B*).

### Differential CJ occupancy between TCR repertoires

From the CJ occupancy matrix, *y*^*k*×*r*^ that represents a the number of T cells in each CJ and repertoire, *ClustIRR* quantifies the relative abundance of cells in each CJ of TCR repertoires, which is also referred to as the CJ *occupancy*. Therefore, we designed a likelihood model *M* for Bayesian inference, which describes the *K*-dimensional CJ occupancy vector *y_i_* for TCR repertoire *i*, as a multinomial model:

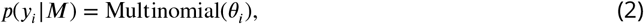

where *θ_i_* is a standard *K*-dimensional simplex CJ occupancy probabilities. Empirically we know that probabilities in *θ_i_* are highly variable between TCR repertoires. To account for this source of variation, the model treats the corresponding coefficients *θ_i_* as random samples drawn from a Dirichlet distribution with parameter vector *ϕ_i_* ∈ ℝ^>0^, where *ϕ_i_* is parametrized as the product of the parameter vector *v_i_*, which describes the expected relative frequency of cells for each CJ of TCR repertoire *i*, and the occupancy parameter, *κ*, which describes the variation in the observed cell counts among the samples (larger values of *κ* induce less between-sample variation from the Dirichlet expectations *v*). This hierarchical modeling framework results in a Dirichlet-multinomial distribution over observed cell counts, which accounts for overdispersion, where the observed variability in cell counts exceeds the level expected under a multinomial model with fixed proportions (*θ_i_*):

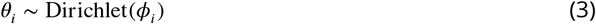

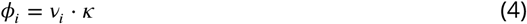

We defined *v_i_* as the softmax function of a vector defined by the linear combination of the intercept parameters, *α* = [*α*_1_, *α*_2_, …, *α*_*k*_], which describe the average (or baseline) cell occupancy in each CJ; and the slope parameter vector, *β*_*ii*1_, *β*_*i*2_, …, *β*_*ik*_], which describes the effect on CJ occupancy of changing from baseline to TCR repertoire *i* for each of the *k* CJs:

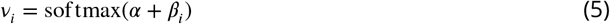

where the softmax function maps *x* ∈ ℝ^*k*^ to the *K*-dimensional simplex by:

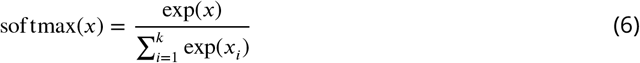

The weakly informative priors assigned to *α*, *β* and *κ* are defined by

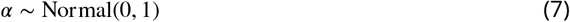

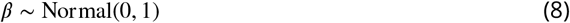

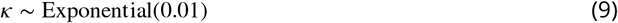

The priors were assessed by prior predictive checks (data not shown). *M* was implemented in Stan (***Carpenter et al., 2017***). Inference of the parameters of *M* was executed with *rstan* (R-package, version 2.32.6) using the No-U-Turn sampler by running a Markov chain Monte Carlo (MCMC) simulation with four chains of 1,750 iterations each, including 750 warm-ups. To test the validity of our model, we performed posterior predictive checks. Furthermore, we inspected the potential scale reduction factor (^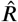^), the effective number of samples in the bulk and tails of the posteriors (ESS-bulk and ESS-tail) and information about divergences during the MCMC sampling to check for convergence. *ClustIRR* outputs the posterior means, medians and the 95% highest density intervals (95% HDIs) of each parameter.

Differential CJ occupancy is analyzed using two related approaches. First, differential CJ occupancy was quantified as

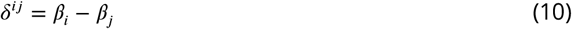

where the parameter vector 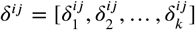 describes the differential CJ occupancy effect size in each CJ. For CJs where 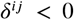 and the 95% HDIs of 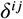 lie mostly or completely below 0 (0 = null effect), we have strong evidence of contracting CJ occupancy in TCR repertoire *i* compared to *j*. On the other hand, for CJs where 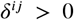 0 and the 95% HDI of 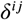 lie mostly or completely above 0, we have strong evidence of expanding CJ occupancy in TCR repertoire *i* compared to *j*. Distributions with the 95% HDIs more or less centered around 0 indicate that there is no evidence for differential CJ occupancy between the two TCR repertoires. Note that unclear evidence is not equivalent to no change of occupancy, because for a CJ with 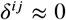 we may also have a wide 95% HDI, including possibilities for positive or negative differential CJ occupancy. Second, differential CJ occupancy was assessed through 95% HDI overlap of *β* coefficients: Non-overlapping HDIs with *β*_*i*_ > *β_j_* indicated strong evidence for contracting occupancy in repertoire j vs. i, while *β*_*i*_< *β_j_* indicated expansion. All cases with overlapping HDIs provided weak evidence for differential CJ occupancy.

### Quantifying Differentially occupant CJs (DCJs) between biological conditions with multiple biological replicates

*ClustIRR* employs a multilevel model, *M_h_*, to quantify convergent differential CJ occupancy between groups of TCR repertoires from *B* biological conditions (vector *b* = {1, 2, …, *B*}). This model extends the repertoire-specific framework (Equation 5) by introducing a group-level hierarchy:

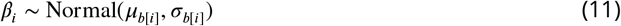

Here, *β*_*i*_ is a *K*-dimensional vector of parameters describing the effects of TCR repertoire *i* on CJ occupancy, where *b*[*_i_*] denotes the biological condition assignment for repertoire *i*. These repertoire-specific coefficients are treated as draws from *K* normal distributions with mean 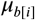_]_and standard deviation 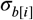_]_. The parameter 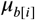 describes the average effect of biological condition *b*[*_i_*] on CJ occupancy across all repertoires in that condition.

This hierarchical structure enables partial pooling of information between TCR repertoires within the same biological condition, leading to improved estimates for each repertoire. The properties of the different populations of parameters are estimated as well, automatically controlling the amount of pooling applied to the TCR repertoire-specific parameters (***Gelman et al., 2012***), with parameters having greater uncertainty pulled more strongly towards population means. The condition-specific means (for condition *j*, where *j* ∈ {1, …, *B*}) are modeled with a weakly informative prior:

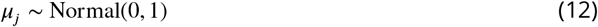

In designs with multiple biological replicates per condition, condition-specific overdispersion parameters *κ_j_* are also inferred, allowing different biological conditions to have distinct levels of variability in CJ occupancy. The weakly informative priors assigned to σ*_j_* and *κ_j_* are defined by:

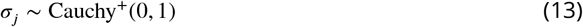

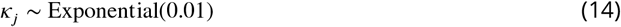

To quantify the differential CJ occupancy between two biological conditions, *x* and *y, ClustIRR* computes an absolute difference between the corresponding *µ* coefficients as described in Equation 10.

### Annotation of T cell clonotype specificity using VDJdb and McPAS-TCR

To annotate T cell clonotypes from our datasets with their putative antigen specificity, we downloaded human and mouse CDR3*α* and CDR3*β* sequences with associated antigen specificity annotations from two public databases: VDJdb (***Shugay et al., 2018***) (https://vdjdb.cdr3.net; accessed June 2024) and McPAS-TCR (***Tickotsky et al., 2017***) (https://friedmanlab.weizmann.ac.il/McPAS-TCR/; accessed January 2025). For annotation, we compared the CDR3*α* and CDR3*β* sequences of TCR clonotypes in our datasets against those in VDJdb and McPAS-TCR. Only identical matches between query (dataset) and reference (database) sequences were retained, and in such cases, the corresponding antigen specificity annotations from the databases were assigned to the query sequences.

### Quantifying TCR repertoire diversity

To quantify the clonotype diversity of TCR repertoires we treated TCR repertoires as generalized communities, in which diversity reflects both richness and evenness, and characterized the TCR clonotype diversity using Normalized Rank Abundance Distributions (NRADs) (***Saeedghalati et al., 2017***). Normalization of the rank-abundance-distribution (RAD) of T cell clonality was done by MaxRank normalization as implemented in R-package *RADanalysis* (version 0.5.5). Here, the MaxRank was set to the minimum dimension of rank abundance vectors for all tested TCR repertoires in a given dataset. Normalized RADs (NRADs) were computed by 1,000-fold averaging, and were visualized with R-package *ggplot2* (version 3.5.1).

### Comparison of relative frequency of DCJs

To analyze the proportion of DCJs in a given T cell receptor (TCR) repertoire from Dataset 2, we employed a hierarchical binomial regression model implemented in Stan. The model accounts for variation in DCJ proportions across four conditions: (1) contracting DCJs in responders; (2) contracting DCJs in non-responders; (3) expanding DCJs in responders; and (4) expanding DCJs in non-responders.

Each observation consists the number of DCJs (*y*) detected within the total number of CJs (*n*) for a given patient. We specified a Bayesian likelihood model to describe these observations for patient *i* ∈ {1, 2, …, 20}, where *y_i_* follows a binomial distribution:

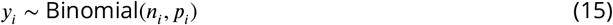

where the probability ***P_i_*** of DCJs is modeled using the inverse logit transformation of a linear predictor that includes an intercept (*α*) and a patient-specific effect (*β*_*i*_):

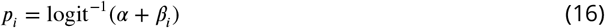

The parameter *α* represents the overall baseline DCJ probability on the inverse logit scale, while *β*_*i*_ captures patient-specific effects. To model variation within groups, we assume a hierarchical structure:

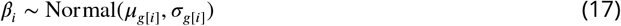

with 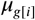 represents the group-level mean effect for condition (group) *g*[*_i_*] ∈ {1, 2, 3, 4}; and 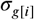 as the within-group standard deviation. We assign weakly informative priors on all parameters:

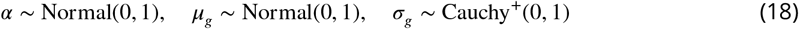

To quantify differences in DCJ proportions between responders and non-responders in contracting (*c*) and expanding (*e*) conditions we estimated the parameters *δ* _*c*_ and *δ _e_*:

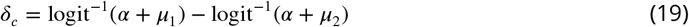

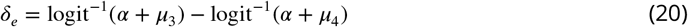

The model was implemented in Stan (***Carpenter et al., 2017***). Inference was performed with *rstan* (R-package, version 2.32.6) using the No-U-Turn sampler by running a Markov chain Monte Carlo (MCMC) simulation with 4 chains of 2,000 iterations each, including 1,000 warm-ups. To test the validity of our model, we performed posterior predictive checks (data not shown), and inspected the potential scale reduction factor (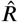), the effective number of samples in the bulk and tails of the posteriors (ESS-bulk and ESS-tail). For each parameter, we report the mean and the 95% highest density interval (95% HDI).

## Software availability

*ClustIRR* (version 1.9.38) is implemented as an R-package, which is freely available as part of the Bioconductor repository.

## Software used to illustrate figures

All figures were generated with *ggplot2* (***Wickham, 2016***) and assembled with *patchwork* (***Pedersen, 2025***).

## Acknowledgments

This work was supported by Deutsche Forschungsgemeinschaft (DFG, German Research Foundation) GRK2762 - project number 450917483 (subprojects M1) to Daniel Hoffmann. We acknowledge support from the Open Access Publication Fund of the University of Duisburg-Essen. We are grateful to Parse Biosciences and ***Formenti et al***. (***2018***) for making their datasets publicly available.

