## Supplementary for "Uncertainty-aware quantitative analysis of the structure and dynamics of T cell receptor repertoires"

### ClustIRR: Uncertainty-aware quantitative analysis of the structure and dynamics of T cell receptor repertoires — <u>Supplementary material</u>

This document includes:
- Supplementary Figures S1–S15
- Supplementary Table S1

#### Supplementary Figures

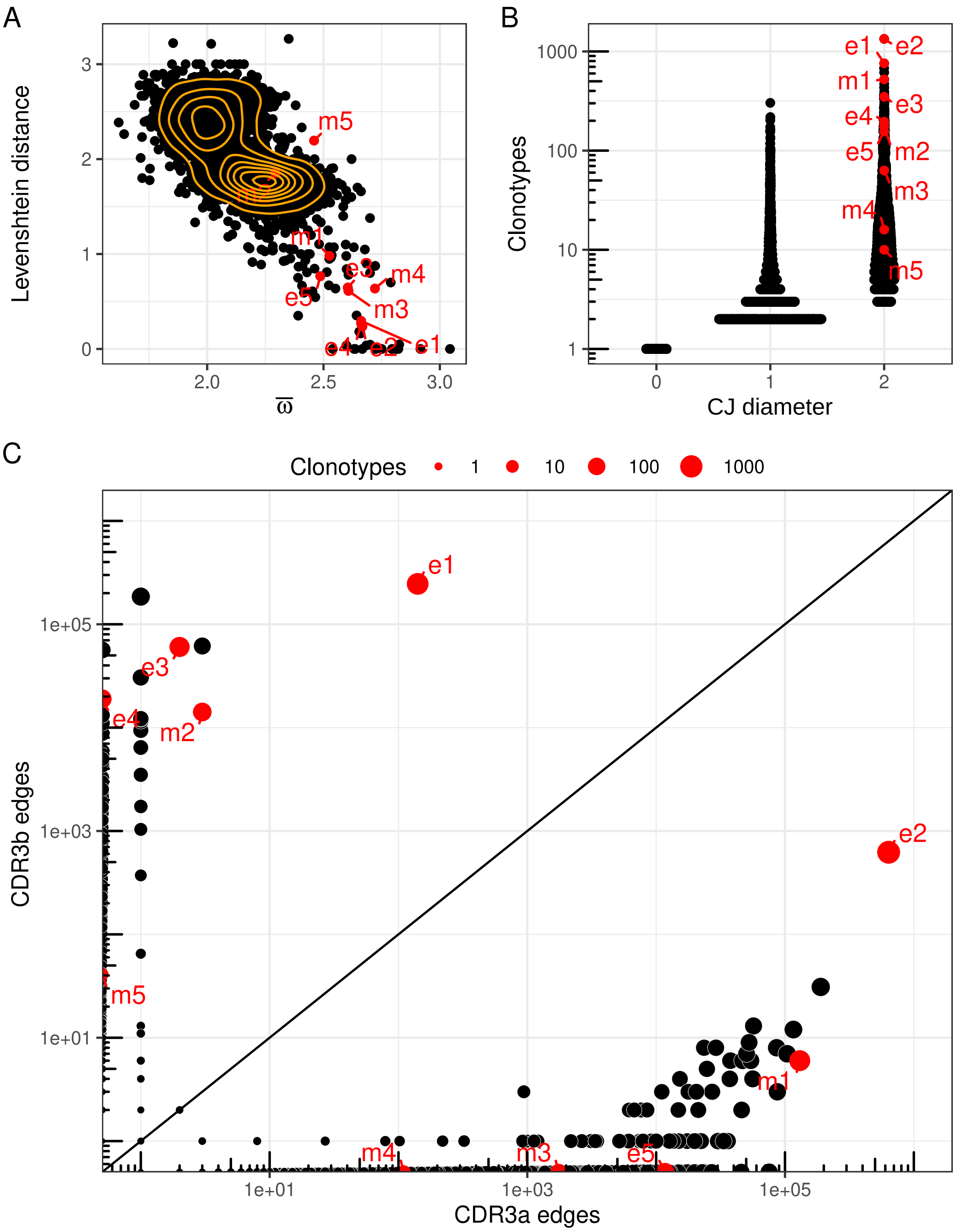


Supplementary Figure S1. Properties of Communities on the Joint graph (CJs). (A) Relationship between mean edge weights ($\bar{\varpi}$, x-axis) and Levenshtein distances (y-axis). (B) Mean unweighted CJ diameter (x-axis) versus CJ size (y-axis). (C) Number of edges driven by CDR3$\alpha$ similarity (x-axis) versus CDR3$\beta$ similarity (y-axis). Five CJ from repertoires E (e1–e5) and M (m1–m5) are labeled.

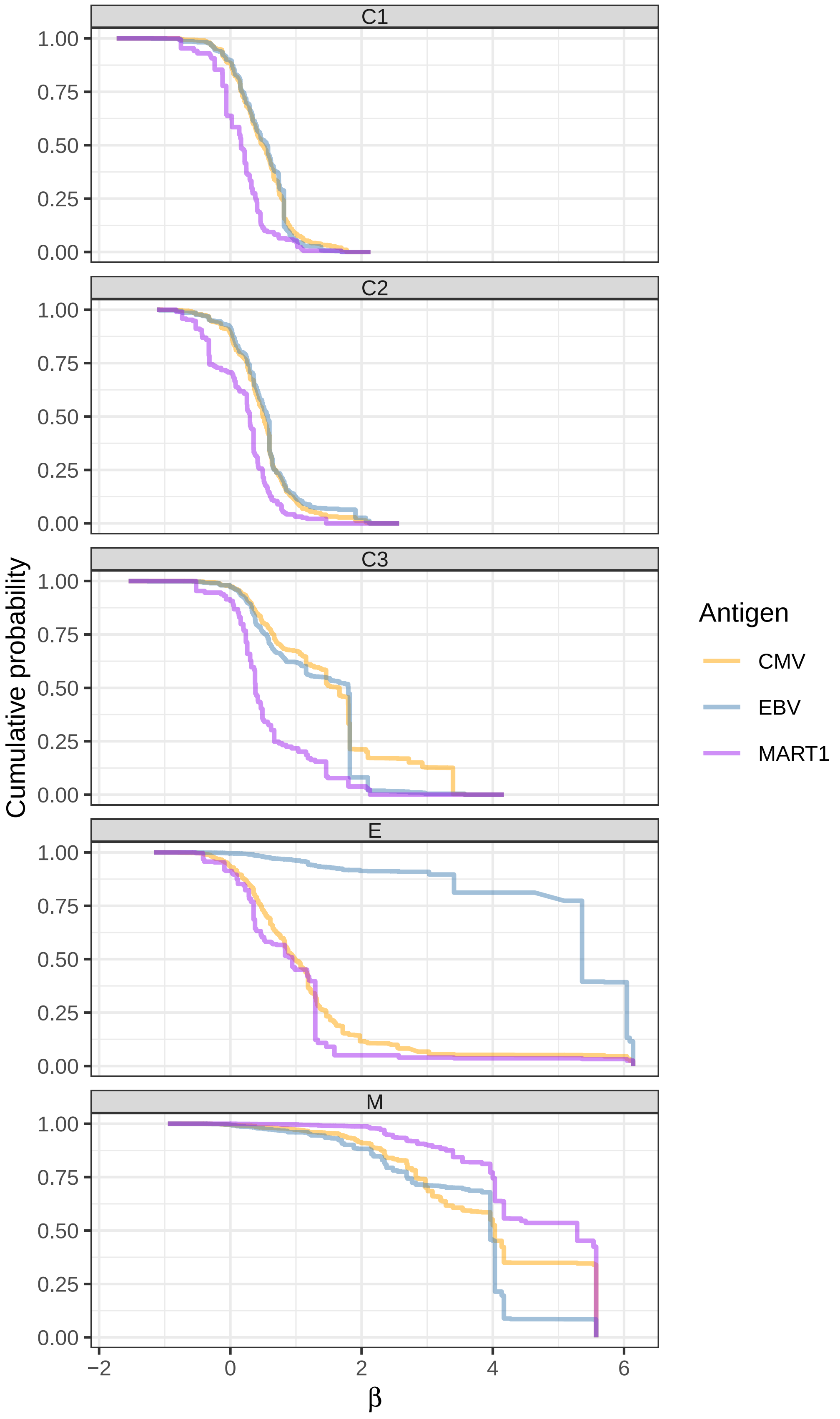


Supplementary Figure S2. Cumulative probability (y-axis) of T-cells specific for antigens (color-code) in CJs sorted based on their $\beta$ coefficients (x-axis). Panels show five TCR repertoires.

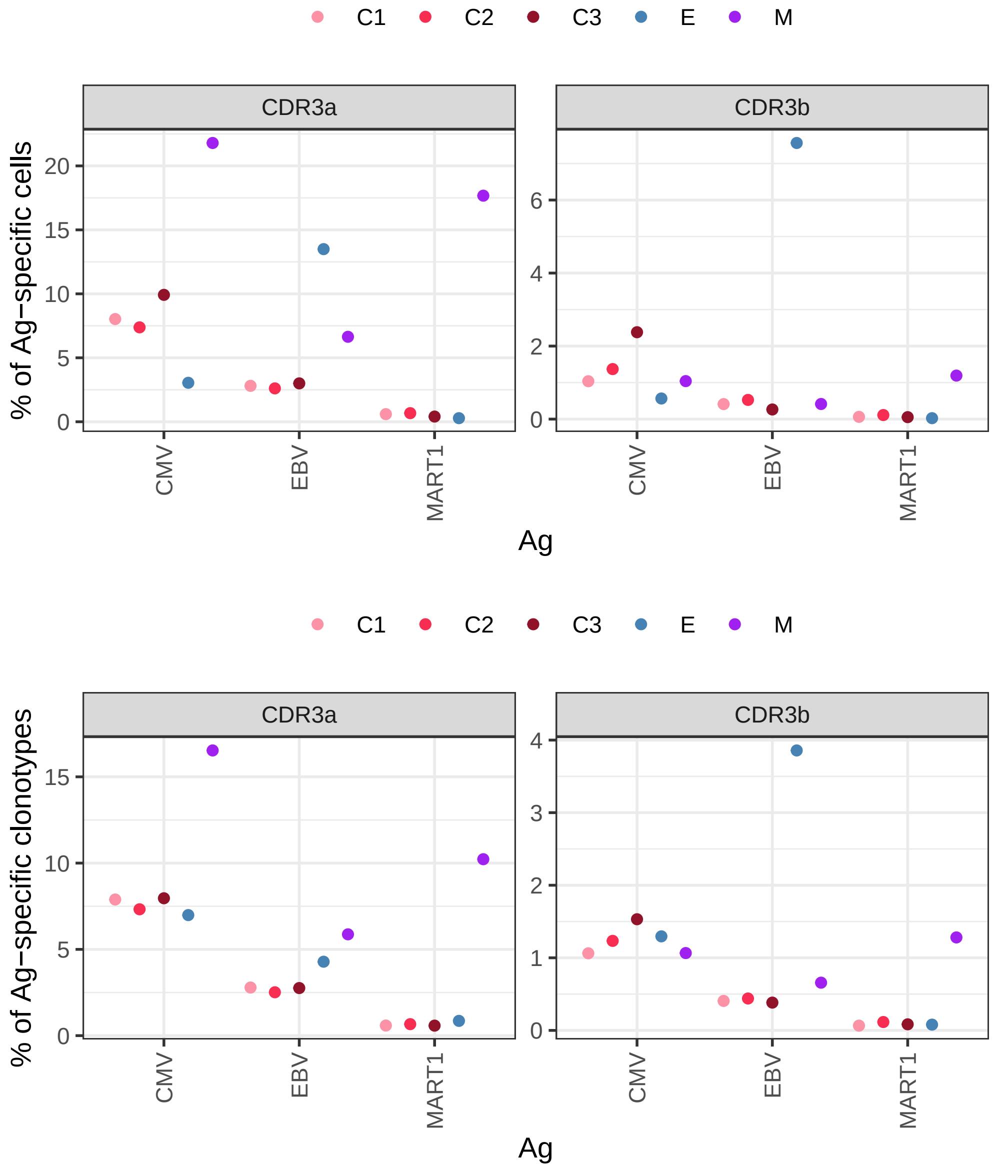


Supplementary Figure S3. Percentage (y-axis) of cells (top row) or clonotypes (bottom row) of T cells specific for EBV-, MART1-, and CMV-derived antigens (x-axis) in each sample (code). The left and right panels show results based on CDR3$\alpha$ and CDR3$\beta$ chains, respectively.

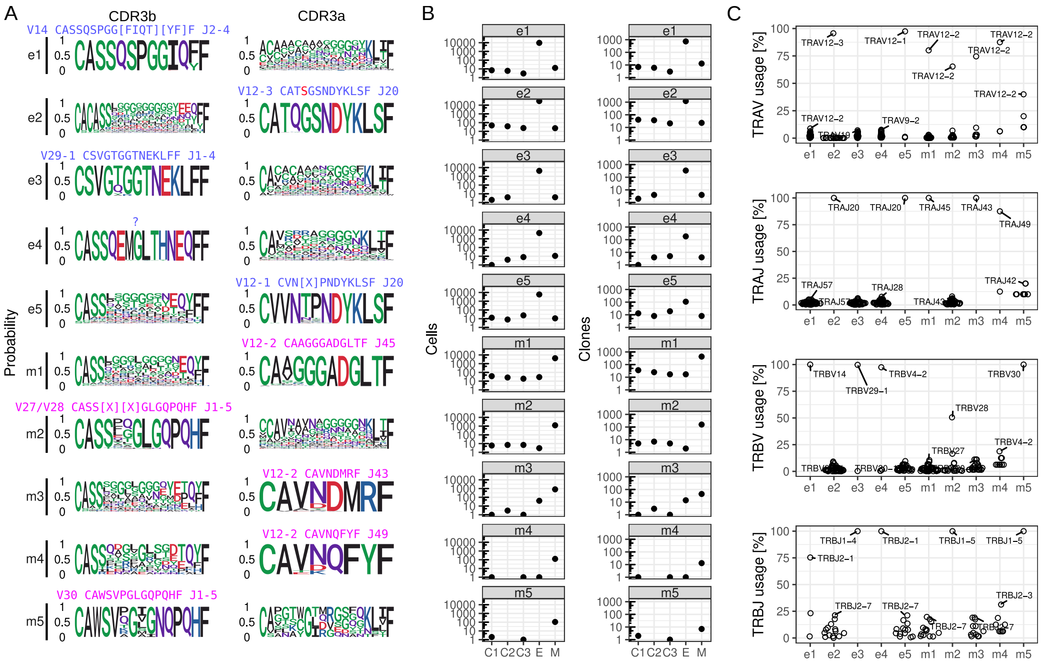

Supplementary Figure S4. (A) Sequence logos of CDR3$\beta$ (left column) and CDR3$\alpha$ (right column) sequences for CJs e1–e5 and m1–m5 (rows). Amino acids are color coded based on their chemical properties. (B) Frequencies of cells (left panel column) and clonotypes (right panel column) from each CJ across the five TCR repertoires, shown in two subpanels. (D) Relative usage of gene segments (TRAV, TRAJ, TRBV and TRBJ) across clonotypes in CJs e1–e5 and m1–m5 (x-axis). Most used gene segments in each CJ are annotated.

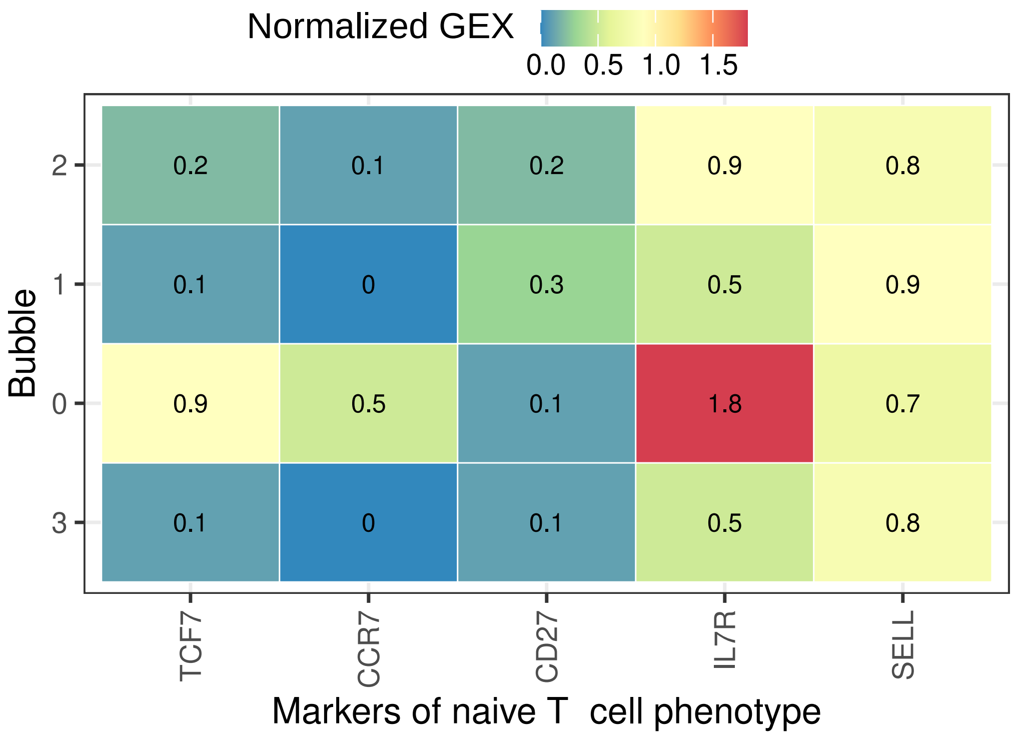


Supplementary Figure S5. Mean gene expression (GEX) of naive T-cell markers (x-axis) across different transcriptional clusters (“bubbles”, y-axis). Color intensity and tile labels indicate the mean expression level.

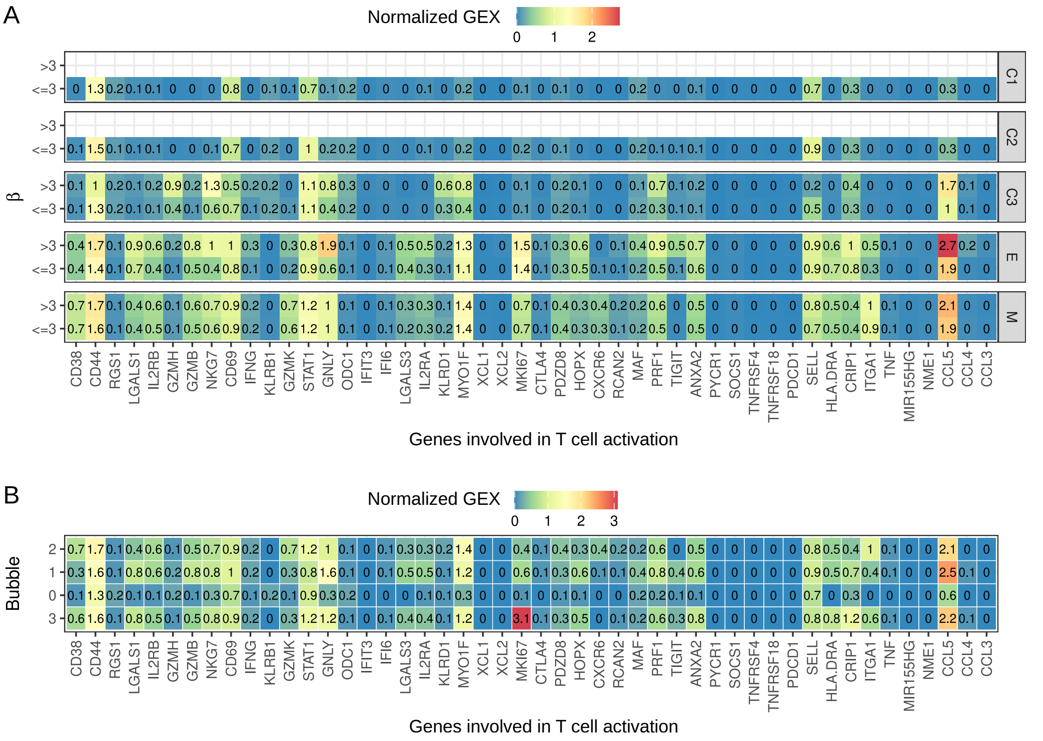


Supplementary Figure S6. Mean gene expression (GEX) of T-cell activation markers across different groupings. (A) Comparison between expanded CJ ($\beta > 3$) and non-expanded CJs ($\beta \leq 3$) across five TCR repertoires (separate panels). (B) Mean expression across transcriptional clusters ("bubbles"). Color intensity and tile labels indicate the mean expression level. Empty tiles denote the absence of cells in the corresponding group.

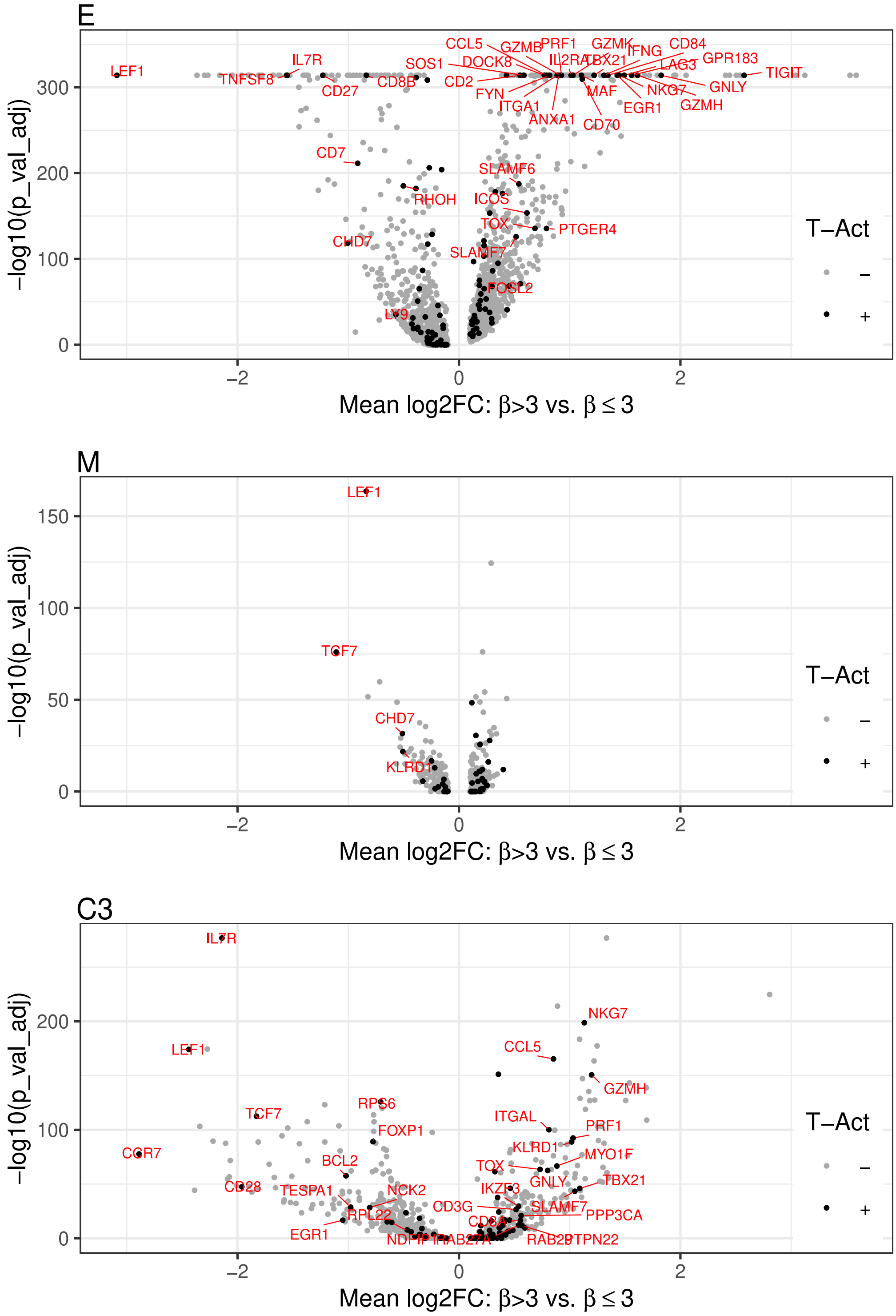


Supplementary Figure S7. Differential gene expression analysis in repertoires E (panel A), M (panel B), and C3 (panel C) comparing CJs with $\beta > 3$ (expanded) versus $\beta \leq 3$. Repertoires C1 and C2 were excluded from this analysis due to a lack of cells in CJs with $\beta > 3$. The x-axis displays the mean $\log_2$ fold change ($\log_2$FC), and the y-axis shows the $-\log_{10}$(adjusted p-value). T-cell activation markers are highlighted as black dots. Markers with $|\log_2 \mathrm{FC}| \geq 0.5$ are labeled in red. Differential expression was performed using the `FindMarkers` function in Seurat with the Wilcoxon Rank Sum test.

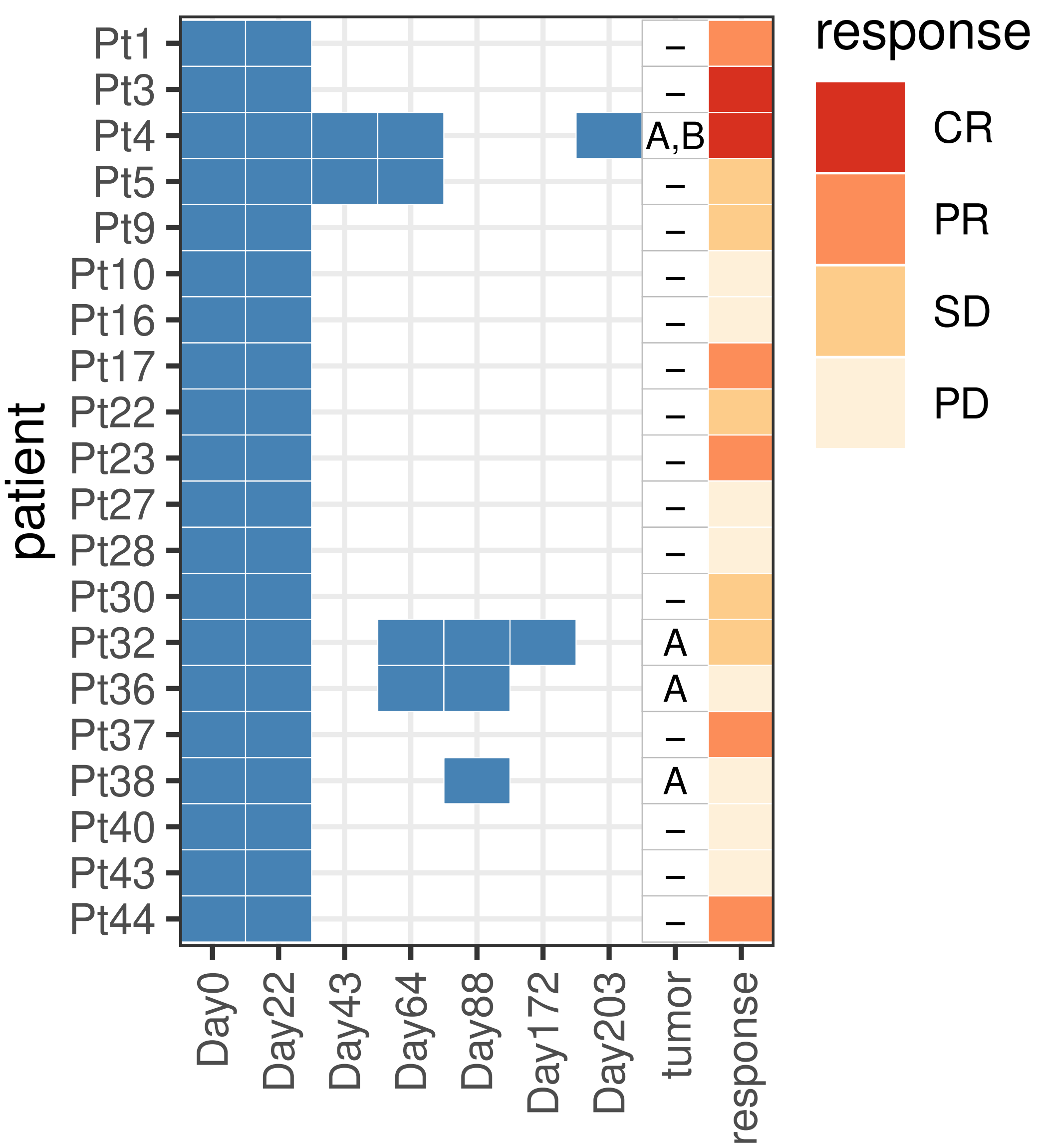


Supplementary Figure S8. Summary of Dataset 2. Blue tiles indicate the time points (x-axis) at which TCR$\beta$ repertoire sequencing was performed on peripheral blood samples for each patient (y-axis). The tile column labeled "tumor" (x-axis) denotes patients for whom TCR$\beta$ repertoire sequencing was performed on archival tumor tissue: Pt4 (two tumors, A and B), Pt32, Pt36, and Pt38. Patient responses to therapy are shown in the final tile column.

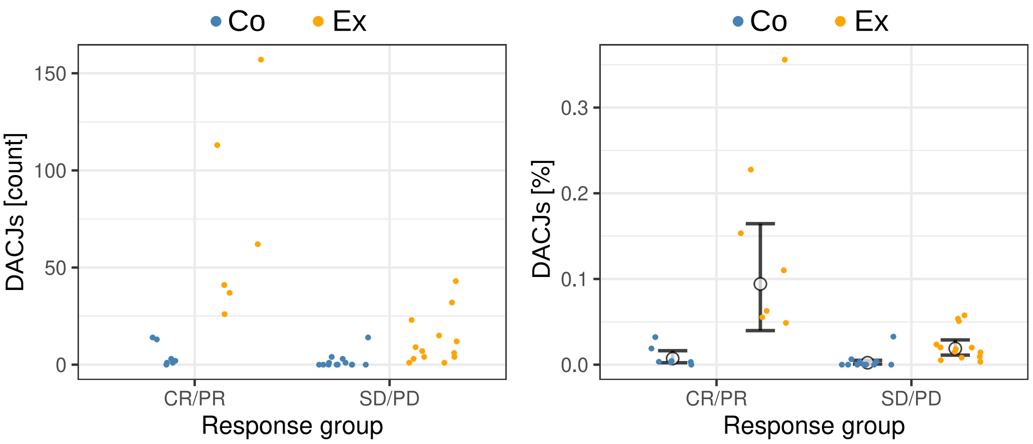


Supplementary Figure S9. Dynamics of Differentially abundant CJs (DCJs) between day 22 and day 0. (A) Count of expanding (E, blue dots) and contracting (C, orange dots) CJs in responders (CR/PR) and non-responders (SD/PD). (B) Percentage of expanding (E, blue dots) and contracting (C, orange dots) CJs in responders (CR/PR) and non-responders (SD/PD), including the overall mean percentages (large black dot) and 95% HDIs (black error bars) in each group. Pairs of groups with statistically different means (95% HDI of absolute difference excluding null effect) are annotated by red horizontal brackets.

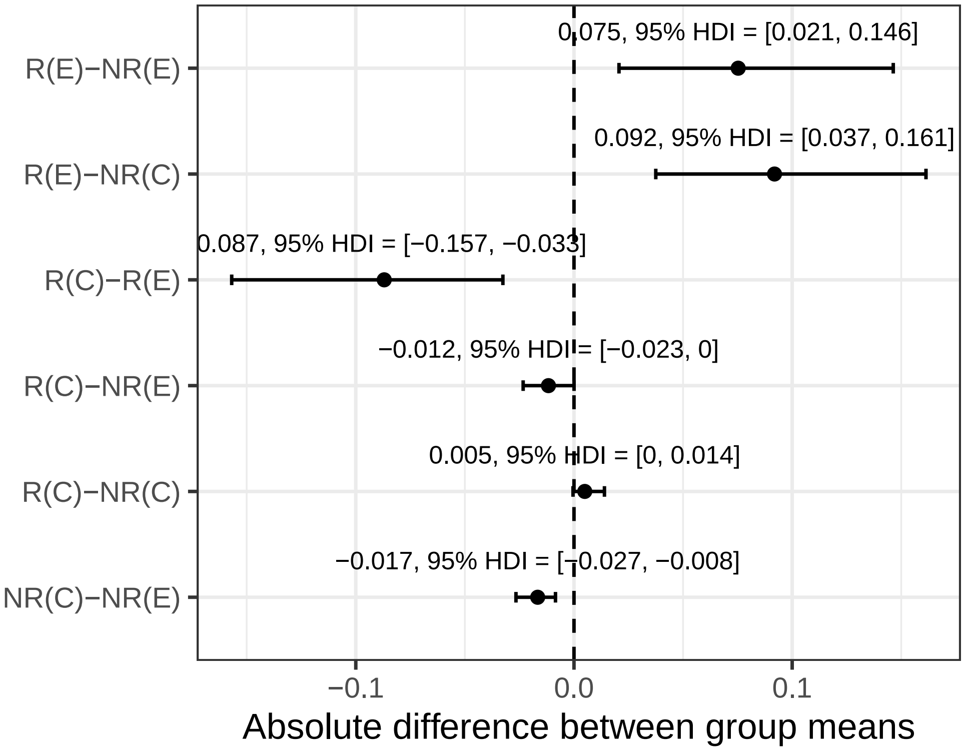


Supplementary Figure S10. Absolute differences in the overall mean prevalence of expanding (E) and contracting (C) Differentially abundant CJs (DCJs) in responders (R) and non-responders (NR). Contrasts are shown on the y-axis. Absolute differences in the means and the corresponding 95% HDIs are shown as dots, error bars, and using labels.

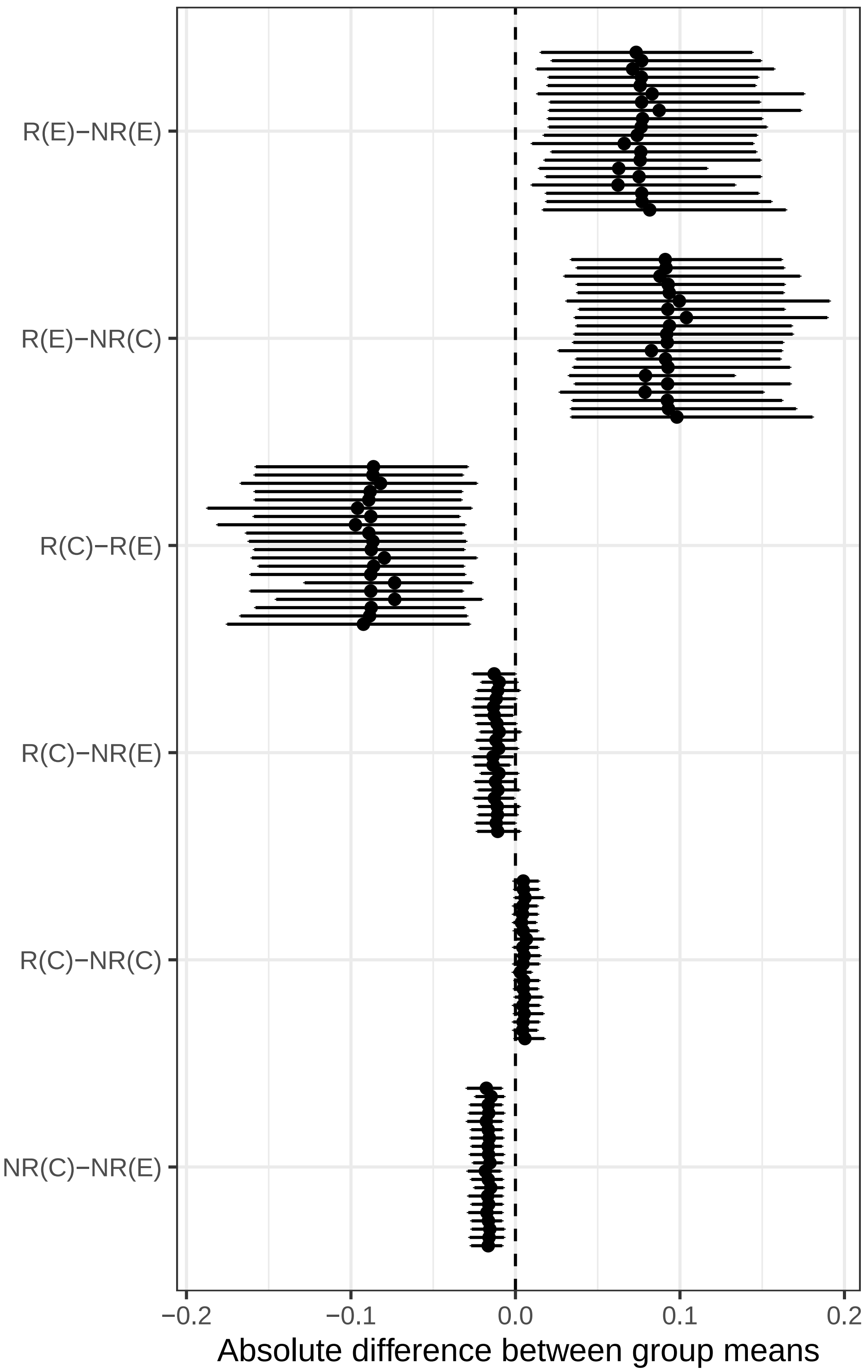


Supplementary Figure S11. Leave one out cross validation (LOOCV) of absolute differences in the overall mean prevalence of expanding (E) and contracting (C) Differentially abundant CJs (DCJs) in responders (R) and non-responders (NR). Contrasts are shown on the y-axis. Absolute differences in the means and the corresponding 95% HDIs are shown as dots, error bars, and using labels. Horizontal jitter was applied to reduce overplotting of overlapping data points from LOOCV analysis.

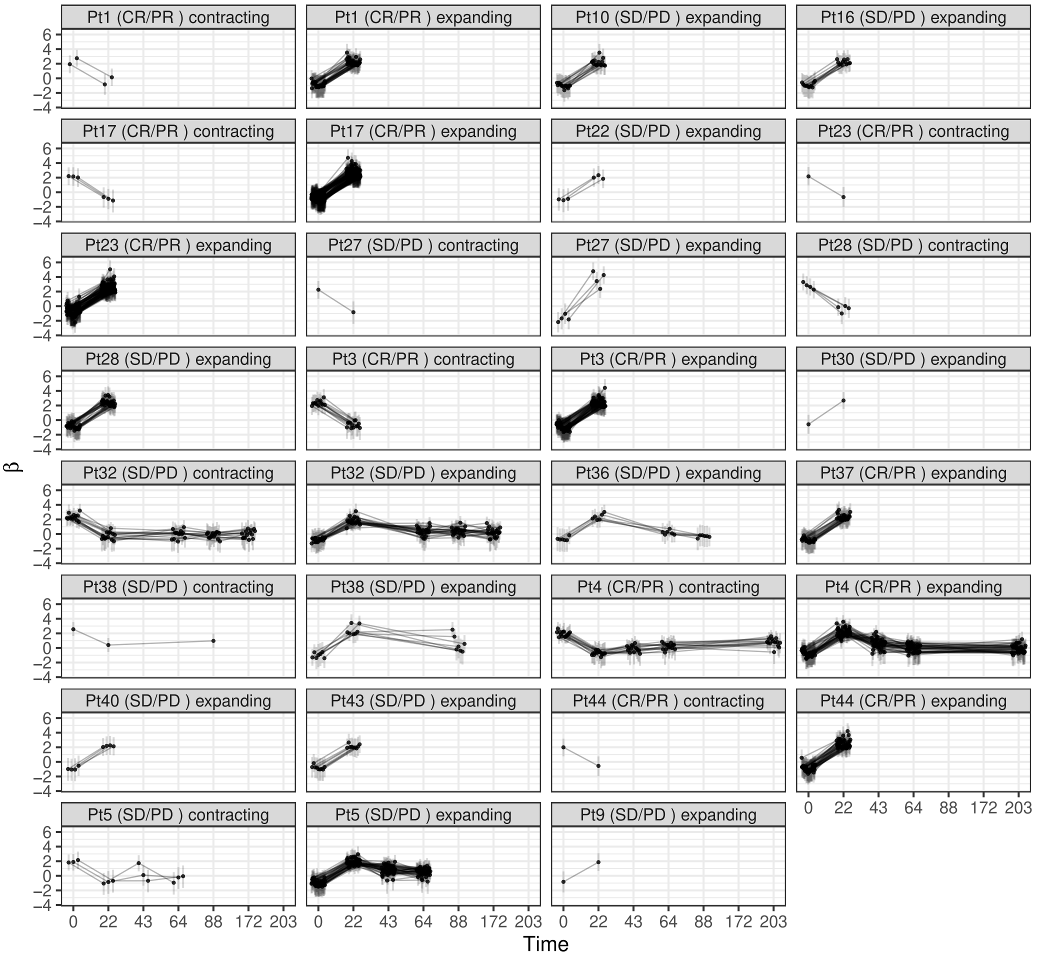


Supplementary Figure S12. Mean CJ intensity ($\beta$, y-axis) and 95% HDI intervals (vertical error bars) for contracting/expanding (panels) DCJs in patients (panel labels) across five timepoints (x-axis).

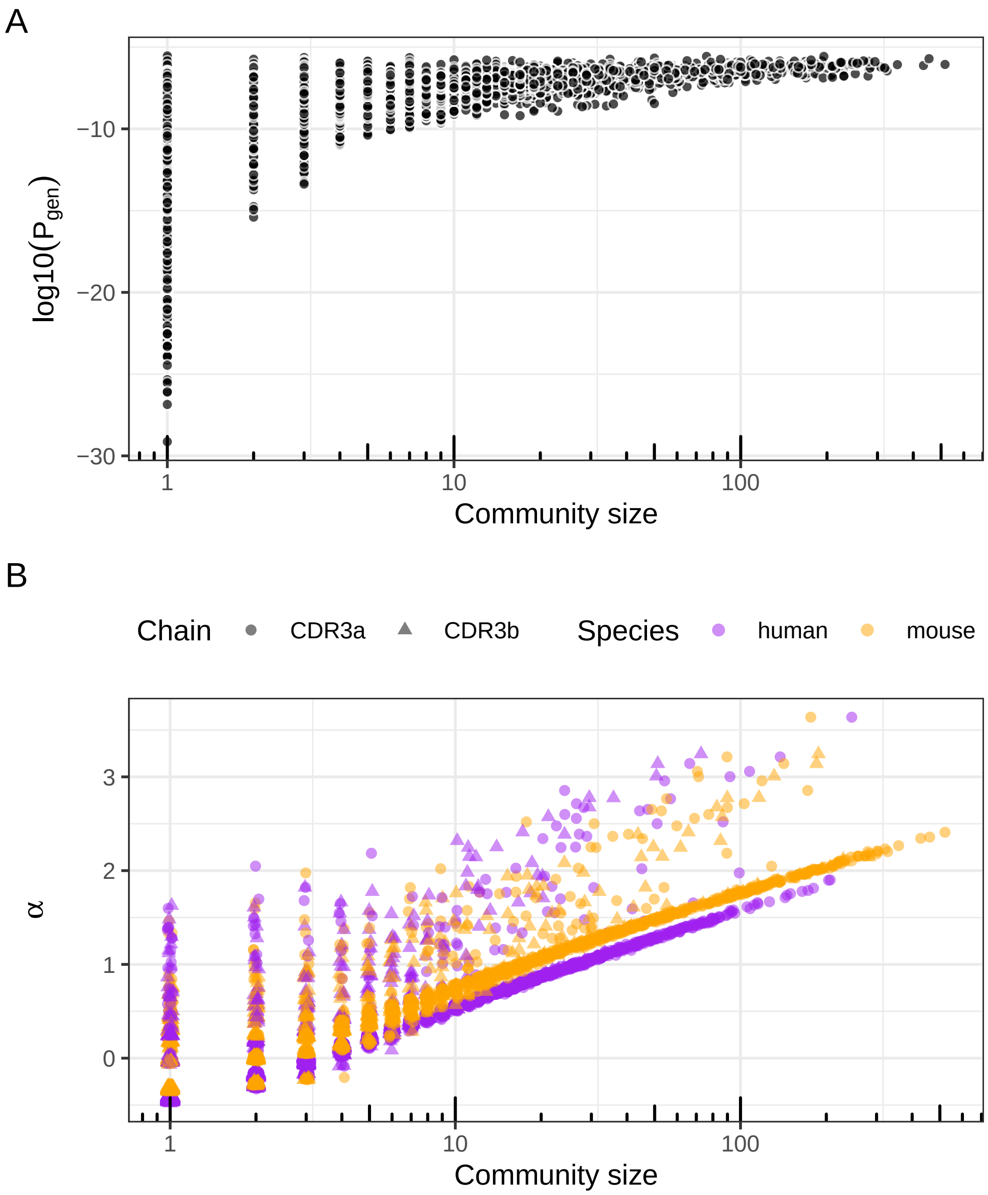


Supplementary Figure S13. Relationship between CJ size and mean generation probability ($P_{gen}$) or model parameter $\alpha$. (A) Mean $\log_{10}(P_{gen})$ of clonotypes within each CJ (y-axis) versus CJ size (x-axis, $\log_{10}$-scaled). (B) Mean baseline CJ occupancy parameter ($\alpha$) (y-axis) versus CJ size (x-axis, $\log_{10}$-scaled). In both panels, each point represents a CJ. In (B), points are colored by species dominance (purple: human; orange: mouse) and shaped by chain type (circles: CDR3$\alpha$; triangles: CDR3$\beta$); and scattered orange and purple points with relatively high $\alpha$ values correspond to CJs containing a mixture of species.

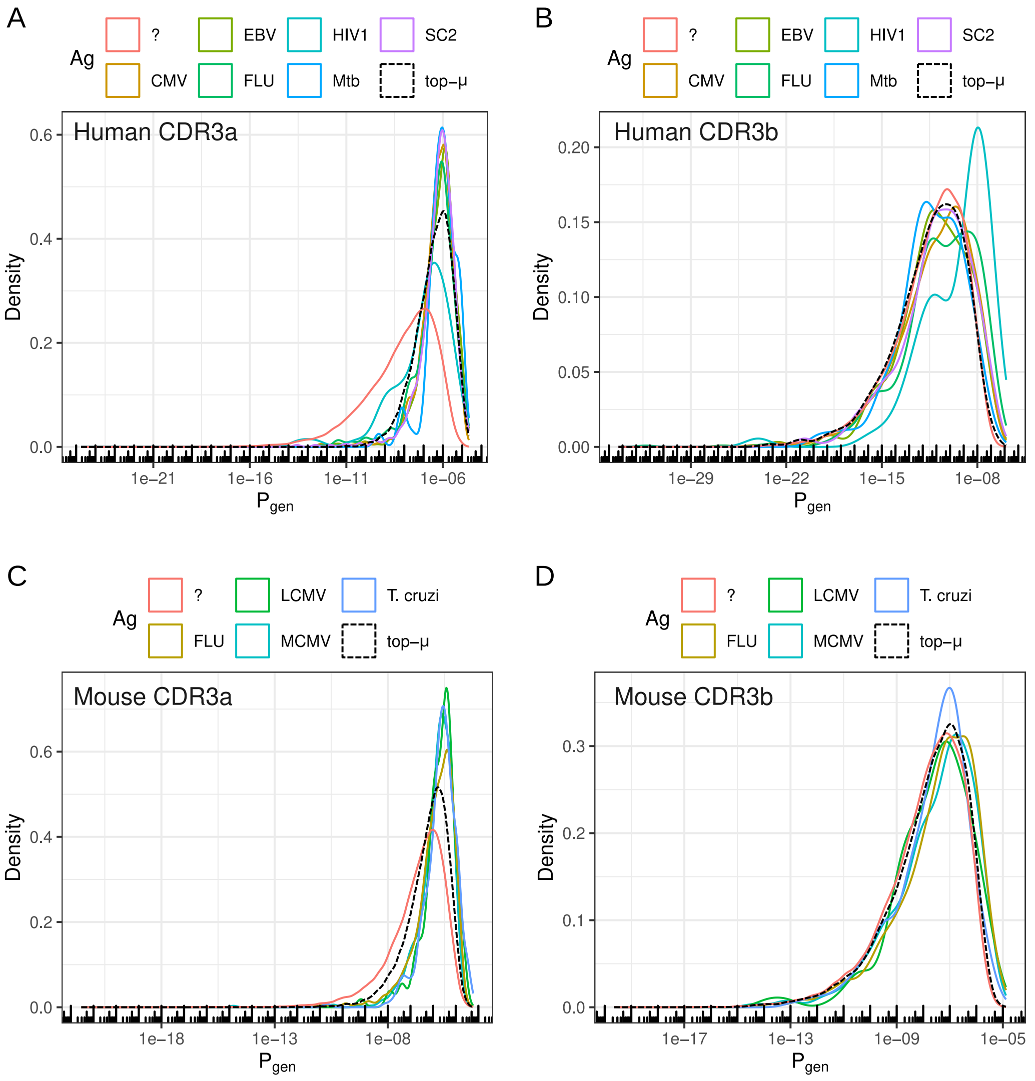


Supplementary Figure S14. Distributions of T-cell receptor generation probabilities ($P_{gen}$) calculated by OLGA using sequences from Dataset 3. (A) Density curves show the distribution of $P_{gen}$ values for human CDR3$\alpha$ sequences across eight groups: six groups specific for pathogens common in humans (CMV, EBV, FLU, HIV-1, Mtb, SARS-CoV-2), one group with unknown antigen specificity (labeled '?'), and one group comprising CDR3$\alpha$s in CJs expanded in humans relative to mice (labeled 'top-$\mu$'). (B) Distribution of $P_{gen}$ values for human CDR3$\beta$ sequences across the same eight specificity groups as in (A). (C) Distribution of $P_{gen}$ values for murine CDR3$\alpha$ sequences across analogous antigen-specific groups (e.g., LCMV, MCMV, FLU, T. cruzi) and control groups. (D) Distribution of $P_{gen}$ values for murine CDR3$\beta$ sequences across the same groups as in (C). Abbreviations: CMV=Cytomegalovirus, EBV=Epstein–Barr virus, FLU=Influenza A virus, HIV-1=Human immunodeficiency virus type 1, LCMV=Lymphocytic choriomeningitis virus, MCMV=Murine cytomegalovirus, Mtb=Mycobacterium tuberculosis, SARS-CoV-2=Severe acute respiratory syndrome coronavirus 2, T. cruzi=Trypanosoma cruzi.

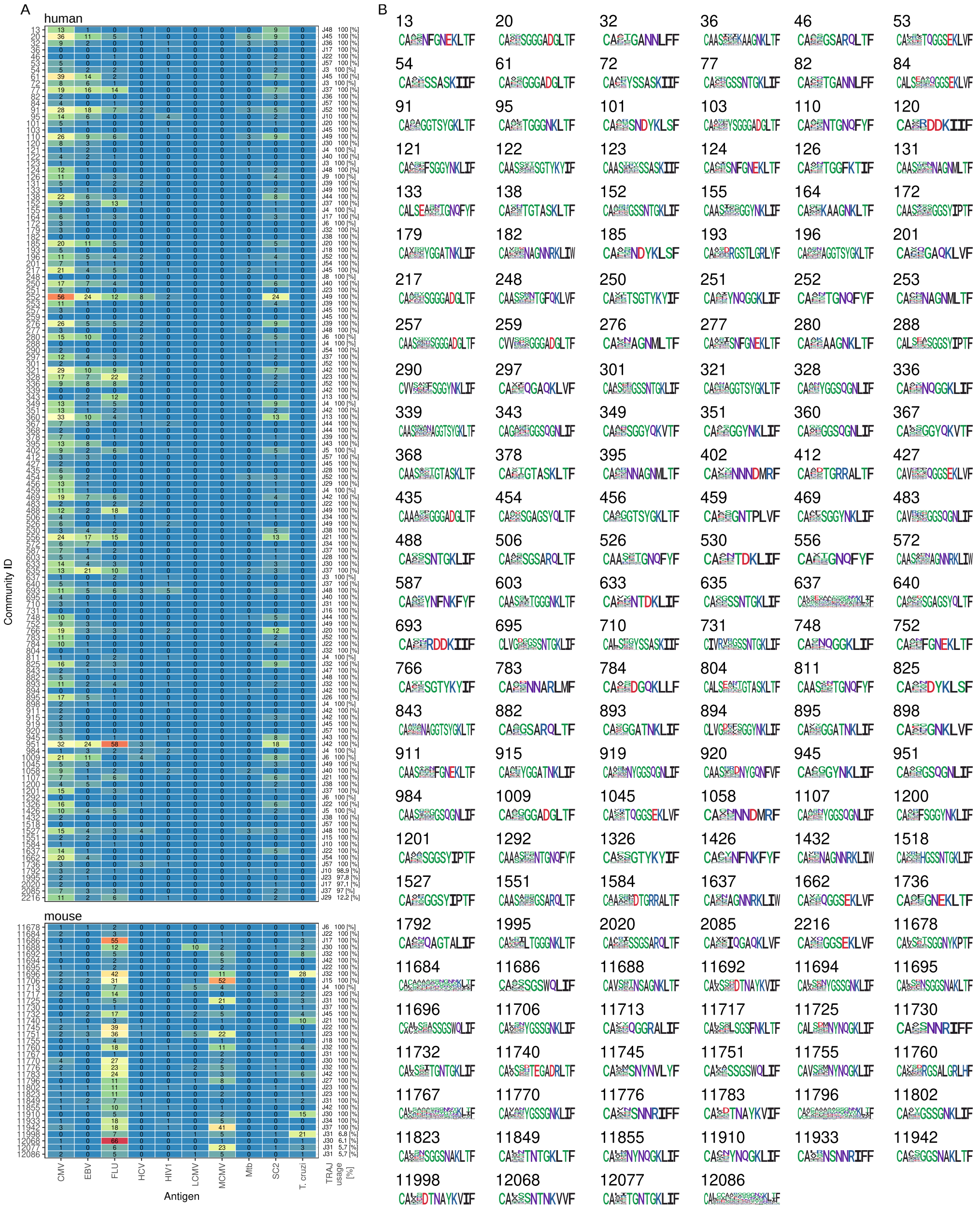

Supplementary Figure S15. (A) Absolute counts of clonotypes specific to ten antigens (x-axis) across each CJ (y-axis: CJ ID) are represented by tile labels and color intensity. The name and relative usage frequency of the predominant TRAJ gene segment are shown in the last column. (B) Sequence logos depict conserved CDR3$\alpha$ motifs for each CJ (labeled by CJ ID). Abbreviations: CMV=Cytomegalovirus, EBV=Epstein-Barr virus, FLU=Influenza A virus, HCV=Hepatitis C virus, HIV-1=Human immunodeficiency virus type 1, LCMV=Lymphocytic choriomeningitis virus, MCMV=Murine cytomegalovirus, Mtb=Mycobacterium tuberculosis, SC2=Severe acute respiratory syndrome coronavirus 2, T. cruzi=Trypanosoma cruzi.

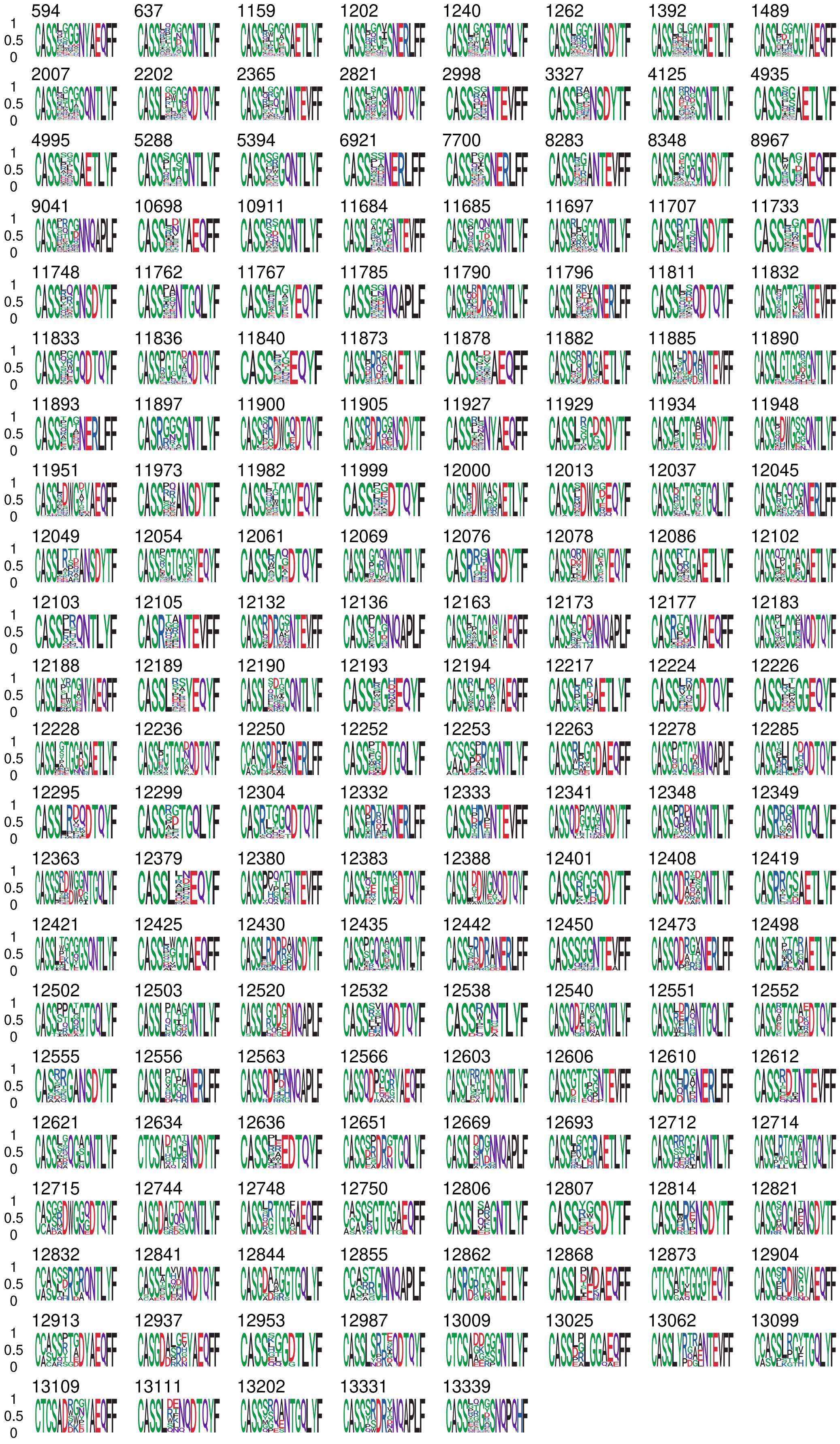


Supplementary Figure S16. Sequence logos depict conserved CDR3$\beta$ motifs for each CJ (labeled by CJ ID).

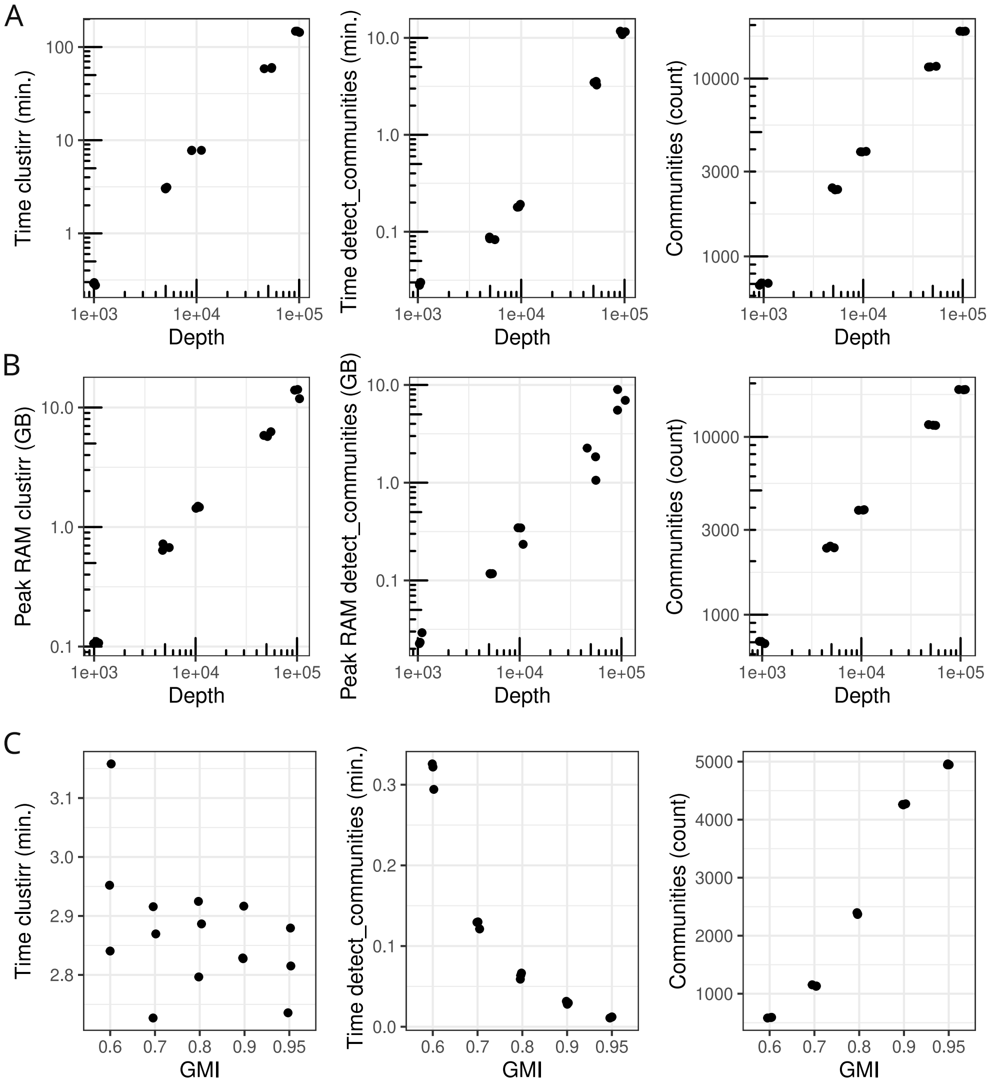


Supplementary Figure S17. Computational performance and parameter sensitivity of ClustIRR. (A) Impact of repertoire depth (number of input paired clonotypes, log10 x-axis) on CPU time, fixed at global minimum identity (GMI) = 0.8. Rows show CPU time for the primary processing function (`clustirr`), CPU time for CJ detection (`detect_communities`), and the number of detected CJs (*N_coms*). (B) Impact of repertoire depth on peak random access memory (RAM, log10 y-axis), fixed at GMI = 0.8. Rows show peak RAM (in gigabytes) for `clustirr`, peak RAM for `detect_communities`, and the number of detected CJs (*N_coms*). (C) Impact of GMI threshold (linear x-axis) on CPU time, fixed at 5,000 paired clonotypes. Rows show CPU time for `clustirr`, CPU time for `detect_communities`, and *N_coms*. All axes in rows B and C are shown on log10 scale. Points represent 3 independent replicates. Results demonstrate that repertoire depth is the primary driver of total runtime and memory, while GMI threshold primarily modulates CJ resolution and clustering speed.

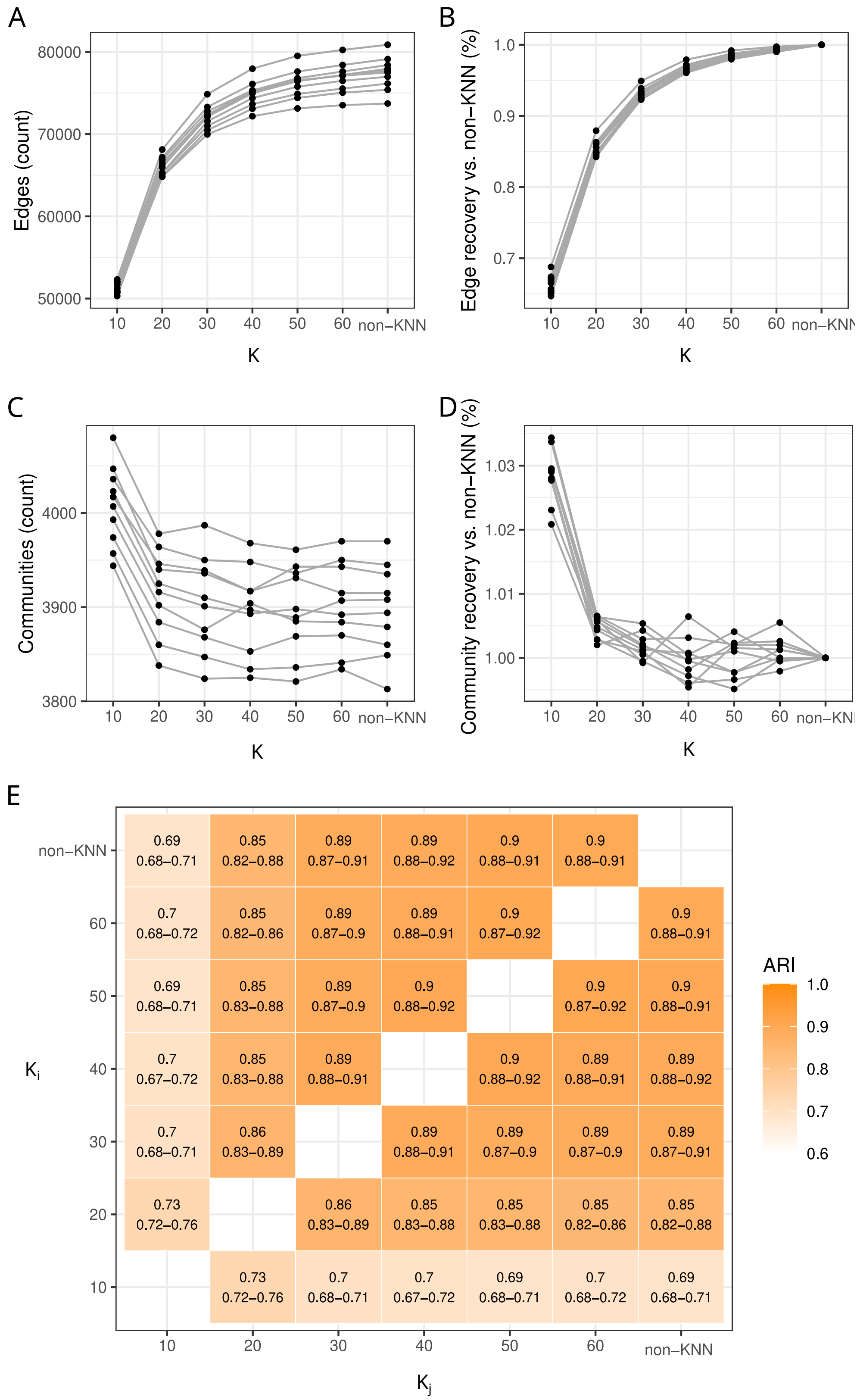


Supplementary Figure S18. Sensitivity of graph topology and clustering stability to KNN parameter $K$. (A) Number of edges and (B) CJs in KNN graphs as a function of $K$ (x-axis), compared against a full graph (non-KNN). (C) Fraction of edges and (D) CJs in KNN graphs relative to the full graph. Lines represent 10 independent replicates with a TCR repertoire depth of 10,000 TCR$\alpha\beta$ clonotypes each. (E) Heatmap tile colors show the mean Adjusted Rand Index (ARI) between clustering assignments for pairs of $K$ values, including the full graph. Diagonal elements (ARI = 1) are omitted. Cell labels display the mean ARI along with the minimum and maximum values across 10 replicates. Results demonstrate the convergence of KNN graph properties toward the full graph as $K$ increases.

### Supplementary Tables

| repertoire | clones | cells |
|---|---|---|
| C1 | 24,384 | 26,035 |
| C2 | 23,431 | 24,377 |
| C3 | 16,715 | 28,076 |
| E | 16,359 | 90,588 |
| M | 8,821 | 46,418 |

Supplementary Table S1. Summary of Dataset 1. TCR repertoire names and the corresponding frequencies of clones and cells. M=MART1, E=EBV.
